# Overcoming donor variability and risks associated with fecal microbiota transplants through bacteriophage-mediated treatments

**DOI:** 10.1101/2023.03.17.532897

**Authors:** Torben Sølbeck Rasmussen, Xiaotian Mao, Sarah Forster, Sabina Birgitte Larsen, Alexandra Von Münchow, Kaare Dyekær Tranæs, Anders Brunse, Frej Larsen, Josue Leonardo Castro Mejia, Signe Adamberg, Axel Kornerup Hansen, Kaarel Adamberg, Camilla Hartmann Friis Hansen, Dennis Sandris Nielsen

## Abstract

**Background:** Fecal microbiota transplantation (FMT) and fecal virome transplantation (FVT, sterile filtrated donor feces) have been effective in treating recurrent *Clostridioides difficile* infections, possibly through bacteriophage-mediated modulation of the gut microbiome. However, challenges like donor variability, costly screening, coupled with concerns over pathogen transfer (incl. eukaryotic viruses) with FMT or FVT hinders their wider clinical application in treating less acute diseases.

**Methods:** To overcome these challenges, we developed methods to broaden FVT’s clinical application while maintaining efficacy and increasing safety. Specifically, we employed the following approaches: 1) Chemostat-fermentation to reproduce the bacteriophage FVT donor component and remove eukaryotic viruses (FVT-ChP), 2) solvent-detergent treatment to inactivate enveloped viruses (FVT-SDT), and 3) pyronin-Y treatment to inhibit RNA-virus replication (FVT-PyT). We assessed the efficacy of these processed FVTs in a *C. difficile* infection mouse model and compared them with untreated FVT (FVT-UnT), FMT, and saline.

**Results:** FVT-SDT, FVT-UnT, and FVT-ChP reduced the incidence of mice reaching the humane endpoint (0/8, 2/7, and 3/8, respectively) compared to the FMT, FVT-PyT, and saline control (5/8, 7/8, and 5/7, respectively) and significantly reduced the load of colonizing *C. difficile* cells and toxin A/B levels. There was a potential elimination of *C. difficile* colonization, with 7 out of 8 mice treated with FVT-SDT testing negative with qPCR. In contrast, all other treatments exhibited the continued presence of *C. difficile*. Moreover, the results were supported by changes in the gut microbiome profiles, cecal cytokine levels and histopathological findings. Assessment of viral engraftment following FMT/FVT treatment and host-phage correlations analysis suggested that transfer of phages likely were an important contributing factor associated with treatment efficacy.

**Conclusions:** This proof-of-concept study show that specific modifications to FVT hold promise in addressing challenges related to donor variability and infection risks. Two strategies lead to treatments significantly limiting *C. difficile* colonization in mice, with solvent/detergent treatment and chemostat-propagation emerging as promising approaches.

## BACKGROUND

During the past decade, it has become evident that various diseases are associated with gut microbiome dysbiosis [1,2], including recurrent *Clostridioides difficile* infections (rCDI) [3,4]. Fecal microbiota transplantation (FMT) from a healthy donor to rCDI patients has proven highly effective in curing the disease, with a success rate exceeding 90% [5,6]. However, FMT faces challenges such as expensive and labor-intensive donor screening [7], donor variability in term of treatment efficacy and reproducibility [7,8], and safety concerns since no screening methods can definitively exclude the transfer of pathogenic microorganisms from the donor. The importance of the latter was highlighted when two patients in the United States experienced severe bacterial infections after FMT, resulting in one fatality [9]. Subsequent safety alerts from the U.S. Food & Drug Administration have warned against potential severe adverse effects associated with the transfer of pathogenic microorganisms during FMT [10,11].

Interestingly, two independent studies [12,13] successfully treated rCDI patients using 0.45 µm sterile filtered donor feces (containing mainly viruses, but possibly also a limited fraction of intact bacteria), a method often referred to as fecal virome transplantation (FVT). The efficacy of FVT was comparable to other clinical studies using FMT (containing bacteria, etc.), suggesting that the gut virome may play an important role when treating rCDI with FMT [12,13]. The gut virome is predominantly composed of bacteriophages (phages), which are host-specific bacterial viruses, but also include eukaryotic and archaeal viruses [14]. The concept of applying FVT from a healthy phenotype to a symptomatic phenotype has been further investigated in preclinical studies. For instance, FVT influenced the composition of the murine gut microbiome following initial perturbation with antibiotics [15]. Additionally, FVT treatment from lean donors alleviated symptoms of metabolic syndrome in three different diet-induced obesity murine models [16–18], and FVT from term piglets prevented the development of necrotizing enterocolitis in a preterm piglet model [19]. FVT has the advantage over FMT in that it significantly diminishes the transfer of viable bacteria, and FVT has recently been demonstrated to be less intrusive for both the gut microbial structure and linked to a reduced likelihood of causing harm to the jejunum in broiler chickens compared to FMT [20]. In addition to the viruses, these FVT preparations would also be expected to contain a certain level of bacterial spores and cells with sizes that allow them to pass through the 0.45 µm filter membrane pores, fecal metabolites, and extracellular vesicles of which contributions to the observed effects following FVT [12,13,15–21] are yet to be elucidated. Furthermore, the sterile filtration process commonly used in FVT does not eliminate the risk of transferring eukaryotic viruses, which previously have been detected in specific pathogen-free mice [22]. While it is possible to screen donor feces for known pathogenic viruses, recent studies have revealed that the human gastrointestinal tract harbors hundreds of eukaryotic viruses with unknown functions [14,23,24]. Most of these viruses are likely harmless to the human host, but it cannot be ruled out that they may contribute to later disease development, as exemplified by the human papillomavirus, which turned out as a significant risk factor for cervical cancer years after infection [25]. Therefore, there are good reasons to minimize the transfer of active eukaryotic viruses when applying FVT to alleviate conditions associated with gut microbiome dysbiosis, particularly when treating individuals with compromised immune systems. In contrast, phages do not actively infect and replicate in eukaryotic cells and are believed to be key players in the successful treatment of gut-related diseases using FVT/FMT [13,16,19,26,27], but the underlying mechanisms are yet poorly understood. FMT and FVT hold the potential to revolutionize treatments for many gut-related diseases, but their widespread use is unlikely due to safety concerns and donor variability. Our objective was, therefore, that we could develop different methodologies that mitigate these challenges while maintaining treatment efficacy. To generate FVTs “free” of eukaryotic viruses, we exploited the fundamental differences in characteristics between eukaryotic viruses and phages. The majority of eukaryotic viruses are enveloped RNA viruses [28,29] and rely on eukaryotic hosts for replication. In contrast, the majority of phages are non-enveloped DNA viruses [28,30] that require bacterial hosts for replication. A solvent/detergent method was applied to inactivate enveloped viruses (FVT-SDT), pyronin-Y was used to inhibit replication of RNA viruses (FVT-PyT), and a chemostat propagated virome (FVT-ChP) was processed to remove the majority of eukaryotic viruses by dilution. Chemostat propagation furthermore has the advantage, that it in principle allows producing more product from the same inoculum, hence increasing reproducibility. These differently processed fecal viromes were evaluated in a C57BL/6J mouse *C. difficile* infection model [31] and compared with a saline solution, FMT (previously shown to effectively treat *C. difficile* infection in preclinical studies [32]), and untreated donor-filtrated feces (FVT-UnT).

This proof-of-concept study represents an important first step towards developing safer and more consistent therapeutic approaches that can effectively target a wide range of gut-related diseases [12,13,16,19,33–35], and potentially supplement FMT with phage-mediated therapies.

## METHODS

### Study design

The *C. difficile* infection model was based on Chen et al. [31] and accommodated the ARRIVE Essential10 guidelines [36]. Forty-eight female C57BL/6J (JAX) mice, 8 weeks old, were obtained from Charles River Laboratories (European distributor of JAX mice) and housed at the AAALAC accredited animal facilities of the Department of Experimental Medicine (AEM, University of Copenhagen, Denmark) in Innovive disposable IVC cages that were replaced once per week. The cages were provided water, food (Altromin 1324 chow diet, Brogaarden, Lynge, Denmark), bedding, cardboard housing, nesting material, felt pad, and biting stem. Upon arrival, the mice were ear tagged, randomly (simple randomization) assigned from the vendor transfer-cages to the IVC cages with 4 mice each and acclimatized for one week (Fig. 1). An antibiotic mixture (kanamycin 0.4 mg/mL, gentamicin 0.035 mg/mL, colistin 850 U/mL, metronidazole 0.215 mg/mL, and vancomycin 0.045 mg/mL) was prepared in the drinking water and provided to the mice through the IVCs for 3 days, and the water was replaced with clean antibiotic-free drinking water for 2 days. Subsequently, the mice received an intraperitoneal injection of clindamycin (2 mg/mL) diluted in sterile 0.9 %(w/v) NaCl water (based on the average body weight of the mice, around 20 g). The mixture and therapeutic doses of antibiotics were performed according to the animal model described by Chen et al. [31]. The aim of the antibiotic treatment was to initiate a gut microbiome dysbiosis that increases the colonization ability of *C. difficile*. A similar sequence of events is also often observed when patients are infected with *C. difficile* during hospitalization after especially prolonged and intense antibiotic treatments [37,38]. Twenty-four hours later, the mice were orally inoculated with 1.21 x 10^4^ CFU of *C. difficile* VPI 10463 (CCUG 19126) via oral gavage. The mice were then divided into six different treatment groups (n = 8): saline (positive control), FMT, FVT-UnT (untreated FVT), FVT-ChP (chemostat propagated virome), FVT-SDT (solvent/detergent treated FVT for inactivating enveloped viruses), FVT-PyT (FVT treated with pyronin-Y for inactivation of RNA viruses). The respective treatments were administered orally by gavage in two doses of 0.15 mL each (FVT solutions were normalized to 2 x 10^9^ Virus-like particles (VLP)/mL), at 18 hours and 72 hours after *C. difficile* inoculation (Fig. 1). The sample size of 8 mice per group was chosen based on previous experiments [16,21] knowingly that the *C. difficile* infection would cause animals being euthanized at different time points, which thereby could challenge the comparability of the different timepoints and affect the statistical power. The reasoning was to accommodate the 3Rs principles (replacement, reduction, and refinement [39]) to reduce the number of animals, due to the severity level of the animal model [31], and that the chosen number of animals were sufficient to assess the potential treatment efficacy of the different processed FVTs. Also, the exclusion of a negative control group (i.e. mice not infected with *C. difficile*) was due to comparable data being available from the original study describing the model [31], again reducing the number of animals needed for the experiment. Treatments and handling of cages (cage 1-12) were performed in the order as Saline, FMT, FVT-UnT, FVT-ChP, FVT-SDT, FVT-PyT (cage 1-6) and repeated with cage 7-12 in same group order. All handling was divided between four authors (TSR, SF, KDT, and AVM) and animal care takers to avoid confounders. All inoculations/treatments by oral gavage were performed blindedly by experienced animal caretakers at AEM. One mouse from the saline treated group (control) was euthanized immediately after oral gavage of *C. difficile*, as the culture was accidentally administered via the trachea, and one mouse from the FVT-UnT group was euthanized 1 week after the 2^nd^ FVT treatment due to malocclusions that had led to malnutrition. This reduced these groups (Saline and FVT-UnT) to n = 7 in both the analysis of survival probability and cytokine profile. Blinded monitoring of the animal health status was performed after *C. difficile* infection to determine if the mice had reached the humane endpoint, leading to immediate euthanization when necessary. The frequency of the health monitoring was adjusted to the current health status of the mice and ranged from every 4^th^ hour (day and night) for the first 3 days after *C. difficile* infection, to every 8-12 hours the following 4 days, and finally every 24 hours per day for the remaining 2 weeks of recovery. The monitoring was supervised blindedly by the study veterinarian (author AVM). The following qualitative parameters were used: physical activity level (i.e., decreased spontaneous or provoked activity), consistency of feces (watery or normal), body posture (hunching or normal), and whether their fur was kept clean or not. The mice were scored on a scale of 0-2: 0 (healthy), 1 (mild symptoms), and 2 (clear symptoms). Mice with a score of 2 that showed no improvement in the above parameters during the subsequent checkup were euthanized. Four authors (TSR, SF, KDT, and AVM) participated in health monitoring, euthanization, tissue and feces sampling. Author TSR was aware of the group allocations at all time points to ensure the correct treatments (but limited participation in health monitoring), while authors SF, KDT, and AVM were blinded at all time points. Fecal pellets were sampled whenever possible at different time points until the mice were euthanized (Fig. 1). Mouse body weights were measured at day 0, 8, 15, 16, 18, 23, 30, and 35 (Fig. S1). At the time of euthanization, samples of the intestinal content from the cecum and colon were taken. A portion of the cecum tissue was fixed in 10 % neutral-buffered formalin (Sarstedt) for histological analysis and stored at room temperature. Another part of the cecum tissue, along with the intestinal content, was preserved for cytokine analysis and stored at -80 °C until use.

**Fig. 1:**
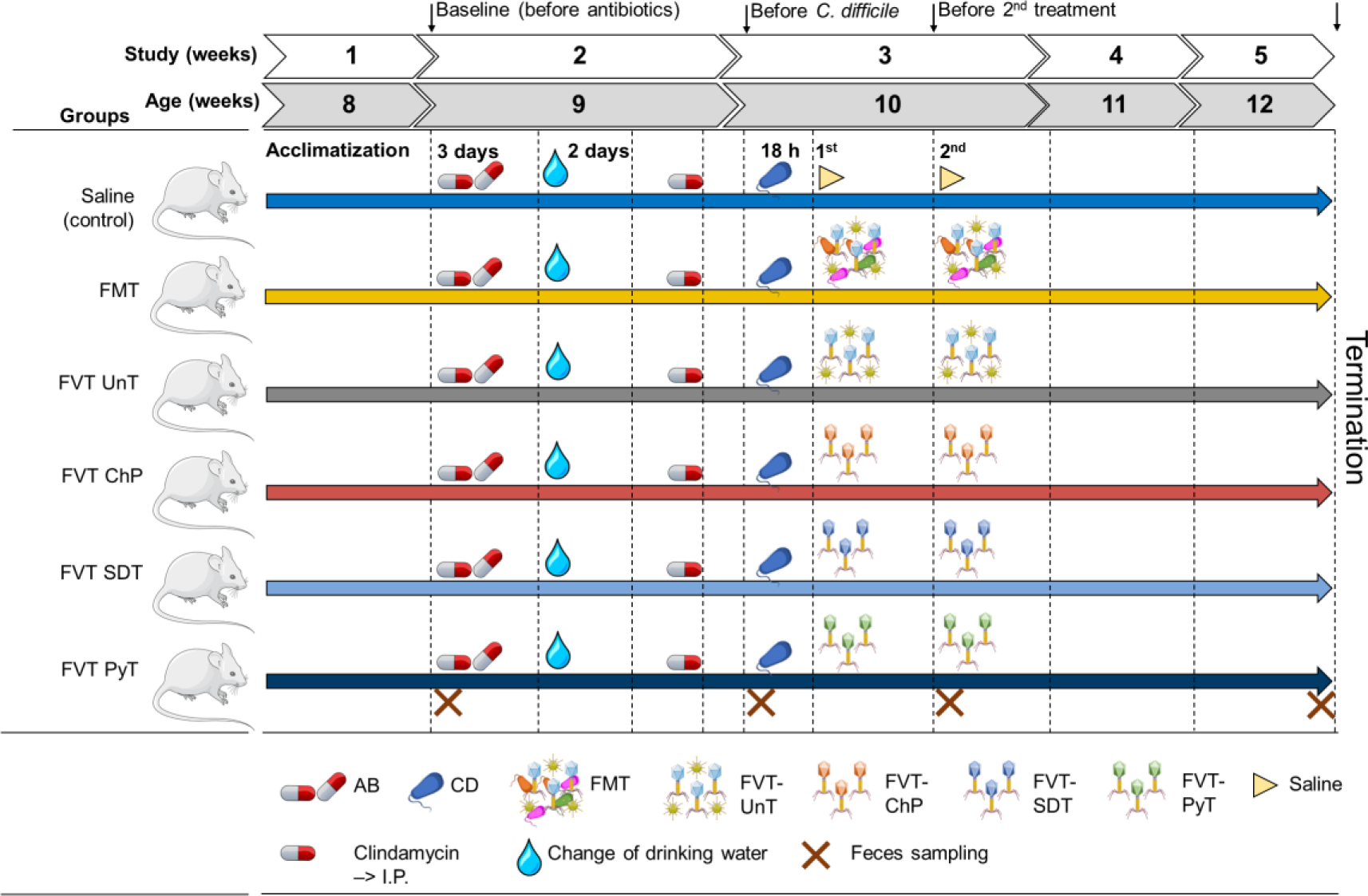
Overview of the animal model. The mice were initially treated with an antibiotic mixture in their drinking water, intraperitoneal (I.P.) injection of clindamycin, and then inoculated with *C. difficile* (∼10^4^ CFU). Eighteen hours after the mice were treated with either saline (as control), FMT (fecal microbiota transplantation), FVT-UnT (fecal virome transplantation – Untreated, i.e. sterile filtered donor feces), FVT-ChP (FVT-chemostat propagated fecal donor virome to remove eukaryotic viruses by dilution), FVT-SDT (FVT-solvent/detergent treated to inactivate enveloped viruses), and FVT-PyT (FVT-pyronin-Y treated to inactivate RNA viruses). Crosses marks time points of feces sampling.

### Clostridioides difficile inoculum

*Clostridioides difficile* VPI 10463 (CCUG 19126), originating from a human infection, was used as infectious agent as in the mouse *C. difficile* infection model described by Chen et al. [31]. The bacteria were cultured in brain-heart-infusion supplement (BHIS) medium [40] with 0.02 %(w/v) 1,4-dithiothreitol (Fisher Scientific) and 0.05 %(w/v) L-cysteine (Fisher Scientific), grown at 37 °C in Hungate tubes (SciQuip), and handled anaerobically as previously described [41]. For solid media, 1.5 %(w/v) agar (Fischer Scientific) was added. Optical density (OD_600nm_) of bacterial cultures was measured with a Genesys30 Visible spectrophotometer (Fischer Scientific). *C. difficile* primarily synthesize its toxins during the stationary phase [42], thus the *C. difficile* culture in its early exponential phase was used as inoculum to minimize transfer of toxins. The *C. difficile* inoculum used for the infection was prepared as follows: a single colony of *C. difficile* was transferred to a Hungate tube containing 10 mL BHIS medium and incubated overnight at 37 °C. Then 150 µL of the *C. difficile* overnight culture was transferred to a new Hungate tube containing 10 mL BHIS medium, incubated for 3.5 hours and its OD_600nm_ was measured. A bacterial calibration curve (OD_600nm_ vs. CFU/mL) of *C. difficile* VPI 10463 was used to dilute the culture to the desired concentration (∼1 x 10^5^ CFU/mL). This constituted the *C. difficile* inoculum. The exact cell concentration of the *C. difficile* inoculum (1.21 x 10^5^ CFU/mL) was evaluated with CFU counts on BHIS agar plates (Fig. S2A-D).

### Host-phage pairs

The bacteria were grown in media and at a temperature suitable for each strain (Table S1). Five phages representing different characteristics (genome, size, and structure), along with their bacterial hosts, were included to assess the influence of pyronin-Y and solvent/detergent treatment on phage activity; *Lactococcus* phage C2 (host: *Lactococcus lactic* DSM 4366), coliphage T4 (host: *Escherichia coli* DSM 613), coliphage phiX174 (host: *E. coli* DSM 13127), coliphage MS2, (host: *E. coli* DSM 5695), and *Pseudomonas* phage phi6 (host: *Pseudomonas* sp. DSM 21482). Solid media were supplemented with 1.5 %(w/v) agar (Thermo Fisher Scientific) for plates and 0.7 %(w/v) for soft agar. An end concentration of 10 mM MgCl_2_ and 10 mM CaCl_2_ was supplemented to media during phage propagation. Plaque activity of phages was evaluated by spot testing where 100 µL bacterial culture was mixed with 4 mL soft agar (temperature at 49 °C), poured to an agar plate, and 10 µL of a phage suspension (dilution series) was deposited on the surface of the solidified soft agar, followed by incubation according to the specific bacterial strain (Table S1).

### Fluorescence microscopy

Virus-like particle (VLP) counts were evaluated of all fecal viromes (FVT-UnT, FVT-SDT, FVT-ChP, and FVT-PyT, Fig. S2E) by epifluorescence microscopy using SYBR Gold staining (Thermo Scientific) as described online dx.doi.org/10.17504/protocols.io.bx6cpraw. The viral concentration was normalized using SM buffer to 2 x 10^9^ VLP/mL per treatment.

### Origin and preparation of intestinal donor material

A total of 54 male C57BL/6N mice were purchased for the purpose of harvesting intestinal content for downstream FVT/FMT applications. Upon arrival, the mice were five weeks old and obtained from three vendors: 18 C57BL/6NTac mice from Taconic (Denmark), 18 C57BL/6NRj mice from Janvier (France), and 18 C57BL/6NCrl mice from Charles River (Germany). We have previously experienced that a high viral diversity can be obtained by mixing the intestinal content from the same mouse strains from three different vendors, and that this approach effectively affected the gut microbiome composition of recipient mice [16,21,43]. The potential importance of high viral diversity on treatment outcome has also previously been suggested in relation to FMT treated *C. difficile* patients [44], and viral diversity have been reported to be positively correlated with donor phage engraftment in a human trial using FMT to treat metabolic syndrome [27]. The mice were earmarked upon arrival, randomly (simple randomization) assigned according to vendor to 3 cages with 6 mice each, and housed at the AAALAC accredited facilities at the Section of Experimental Animal Models, University of Copenhagen, Denmark, following previously described conditions [43]. They were provided with *ad libitum* access to a low-fat diet (LF, Research Diets D12450J) for a period of 13 weeks until they reached 18 weeks of age, which was the planned termination point. Unfortunately, malocclusions resulted in malnutrition for two C57BL/6NRj mice, and they were euthanized before the intended termination date. All mice were euthanized by cervical dislocation, and samples of intestinal content (not feces pellets) from the cecum and colon were collected and suspended in 500 µL of autoclaved anoxic PBS-buffer (137 mM NaCl, 2.7 mM KCl, 10 mM Na_2_HPO_4_, 1.8 mM KH_2_PO_4_). Subsequently, all samples were stored at -80 °C. In order to preserve the viability of strict anaerobic bacteria, 6 mice from each vendor (a total of 18 mice) were sacrificed and immediately transferred to an anaerobic chamber (Coy Laboratory) containing an atmosphere of approximately 93 % N_2_, 2 % H_2_, and 5 % CO_2_, maintained at room temperature. The samples collected from these mice within the anaerobic chamber were used for FMT and anaerobic chemostat cultivation to produce the FVT-ChP. The intestinal content from the remaining 34 mice was sampled under aerobic conditions and used to generate the fecal virome for downstream processing of the FVT-UnT, FVT-SDT, and FVT-PyT treatments. A flow diagram illustrating the aforementioned processes is provided (Fig. S3). An anaerobic growth test was performed for all FVT inoculums to evaluate the level of residual viable bacterial cells or spores (Table S2). It was conducted by spreading 50 µL of undiluted FVT on non-selective Gifu Anaerobe Medium (GAM, Himedia) 1.5 %(w/v) agar plates inside an anaerobic chamber. Two replicates of each FVT were incubated at 37 °C in an anaerobic jar containing an anaerobic sachet (AnaeroGen, Thermo Fisher Scientific) outside the anaerobic chamber for 14 days before CFU counting was performed.

#### Untreated fecal virome (FVT-UnT)

For processing FVT solutions [16], thawed intestinal content from the cecum and colon was suspended in 29 mL autoclaved SM-buffer (100 mM NaCl, 8 mM MgSO_4_·7H_2_O, 50 mM Tris-HCl with pH 7.5), followed by homogenization in BagPage+ 100 mL filter bags (Interscience) with a laboratory blender (Seward) at maximum speed for 120 seconds. The filtered and homogenized suspension was subsequently centrifuged using a centrifuge 5920R (Eppendorf) at 4,500 x g for 30 minutes at 4 °C. The fecal supernatant was sampled for further processing of FVT solutions, while the pellet was resuspended in PBS buffer for bacterial DNA extraction. The fecal supernatant was filtered through a 0.45 µm Minisart High Flow PES syringe filter (Sartorius) to remove bacteria and other larger particles. This step of the FVT preparation does not definitively exclude that extracellular vesicles, bacterial cells, and spores can pass through the 0.45 µm filters. Ultrafiltration was performed to concentrate the fecal filtrate using Centriprep Ultracel YM-30K units (Millipore) that by its design constitute of an inner and outer tube. The permeate in the inner tube was discarded several times during centrifugation at 1,500 x g at 20 °C until approximately 0.5 mL was left in the outer tube, which at this point was considered as a fecal virome. The 30 kDa filter from the Centriprep Ultracel YM-30K units was removed with a sterile scalpel and added to the fecal virome to allow viral particles to diffuse overnight at 4 °C. In order to trace back the origin of specific bacterial or viral taxa, the fecal viromes were mixed based on cages, taking into account the coprophagic behavior of mice [45]. The ultrafiltration of FVT-UnT, -ChP, -SDT, and -PyT is expected to remove the vast majority of metabolites below 30 kDa [46,47]. These fecal viromes were mixed into one final mixture from mice of all three vendors representing the “untreated fecal virome”, FVT-UnT, which was immediately stored at -80 °C. The remaining fecal viromes were stored at 4 °C prior to downstream processing to inactivate the eukaryotic viruses in the fecal viromes by either dissolving the lipid membrane of enveloped viruses with solvent/detergent treatment or inhibit replication of RNA viruses with pyronin-Y.

#### Solvent/detergent treated fecal virome (FVT-SDT)

The solvent/detergent treatment is commonly used to inactivate enveloped viruses, as most eukaryotic viruses possess an envelope, while non-enveloped viruses, including phages, are not affected by this treatment [48,49]. Following the guidelines set by the World Health Organization (WHO) [50] and Horowitz et al. [48] for the clinical use of solvent/detergent-treated plasma, the fecal viromes were subjected to incubation in a solution containing 1 %(w/v) tri(n-butyl) phosphate (TnBP) and 1 %(w/v) Triton X-100 at 30 °C for 4 hours. It is important to note that the majority of inactivation typically occurs within the first 30-60 minutes of the solvent/detergent treatment [50]. The removal of TnBP and Triton X-100 was performed according to the method described by Treščec et al. [51]. In brief, the applied volume of Amberlite XAD-7 in the column was set to 150 % of the theoretical binding capacity to ensure a sufficient removal of TnBP and Triton X-100. The resin column was equilibrated with 0.01 M phosphate buffer (Na_2_HPO_4_ and NaH_2_PO_4_) pH 7.1 containing 0.5 M NaCl until OD_280nm_ was < 0.02. Each solvent/detergent treated fecal virome (mixed by cage) was added separately to the column and OD_280nm_ was measured to follow the concentration of proteins (expected viral particles and other metabolites > 30 kDa) and until OD_280nm_ was < 0.02. A 0.01 M phosphate buffer containing 1 M NaCl was used to release potential residual particles from the resin [51]. The removal of the solvent/detergent agents from the fecal viromes yielded approx. 100 mL viral-flow-through from the column which was concentrated to 0.5 mL using Centriprep Ultracel YM-30K units as described in the previous section. The final product constituted the FVT-SDT treatment and was stored at -80 °C until use.

#### Pyronin-Y treated fecal virome (FVT-PYT)

Pyronin-Y (Merck) is a strong red-colored fluorescent compound. It has been reported to exhibit efficient binding to single-stranded and double-stranded RNA (ss/dsRNA), while its binding to single-stranded and double-stranded DNA (ss/dsDNA) is less effective [52,53]. Initial screening was conducted to determine the optimal conditions for viral inactivation using various concentrations of pyronin-Y, different incubation times, and temperatures for RNA and DNA phages. The fecal filtrate was treated with 100 µM pyronin-Y and incubated at 40 °C overnight to inactivate viral particles containing RNA genomes. To remove the pyronin-Y molecules that were not bound to particles, the pyronin-Y-treated fecal filtrate suspensions were diluted in 50 mL SM-buffer and subsequently concentrated to 0.5 mL using ultrafiltration with Centriprep Ultracel YM-30K units. This process was repeated three times, resulting in a transparent appearance of the pyronin-Y treated fecal filtrate, which constituted the FVT-PyT treatment and was stored at -80 °C until use.

#### Fecal microbiota transplantation (FMT)

The mouse intestinal content that was sampled anoxically (Fig. S3) was diluted 1:20 in an anoxic cryoprotectant consisting of PBS-buffer and 20 %(v/v) glycerol and stored at -80 °C until administration.

#### Chemostat propagated fecal virome (FVT-ChP)

The preparation of the chemostat propagated virome was performed as described previously [54]. Briefly, anaerobic handled mouse cecum content was utilized for chemostat propagation. The culture medium was formulated to resemble the low-fat (LF) diet (Research Diets D12450J) provided to the donor mice as their feed (Table S3), and growth conditions such as temperature (37 °C) and pH (6.4) were set to simulate the environmental conditions present in the mouse cecum. The end cultures, which underwent fermentation with a slow dilution rate (0.05 volumes per hour), exhibited a microbial composition that resembled the initial microbial composition profile of the donor [54]. These batches were combined to form the FVT-ChP treatment and were stored at -80 °C until use.

### Cytokine analysis

Pre-weighted cecum tissue was homogenized in 400 μL lysis-buffer (stock solution: 10 mL Tris lysis-buffer, 100 μL phosphatase inhibitor 1, 100 μL phosphatase inhibitor 2, and 200 μl protease inhibitor) (MSD inhibitor pack, Meso Scale Discovery) using a FastPrep Bead Beater Homogenizer (MP Biomedicals), and centrifuged (8,000 x g; 4 °C; 5 minutes). Samples were diluted 1:2 and analyzed for IFN-γ, GM-CSF, IL-15, IL-6, IL-10, KC/GRO, MIP-2, TNF-α IL-17A/F, and IL-22 in a customized metabolic group 1 U-PLEX (MSD) according to manufacturer’s instructions. Samples were analyzed using the MESO QuickPlex SQ 120 instrument (Meso Scale Discovery) and concentrations were extrapolated from a standard curve using Discovery Workbench v.4.0 (Meso Scale Discovery) software. Measurements out of detection range were assigned the value of lower (set to 0) or upper detection limit. The cytokine analysis was performed by a blinded investigator.

### Histology and cytotoxicity assay

Formalin-fixed, paraffin-embedded cecum tissue sections were stained with hematoxylin and eosin for histopathological evaluation by a blinded investigator (author AB). A composite score was assigned, taking into account the following pathological features: 1) immune cell infiltration, 2) submucosal edema or hemorrhage, 3) epithelial injury, each with a range of severity/extent as follows: 0: none, 1: mild, 2: moderate, 3: severe) for a cumulative pathology grade between 0 and 9 [16]. Cecum tissue samples with mechanical damage were excluded for the analysis.

The RIDASCREEN *C. difficile* Toxin A/B ELISA kit (r-biopharm) was used to measure the toxin concentrations in the mice feces by following the instructions of the manufacturer. The OD_450nm_ was measured with a Varioskan Flash plate reader (Thermo Fisher Scientific).

### qPCR measuring *C. difficile* abundance

*C. difficile* in the fecal samples was enumerated using quantitative real-time polymerase chain reaction (qPCR) with species-specific primers (C.Diff_ToxA_Fwd: 5’-TCT ACC ACT GAA GCA TTA C-3’, C.Diff_ToxA_Rev: 5’-TAG GTA CTG TAG GTT TAT TG-3’ [55]) purchased from Integrated DNA Technologies. Standard curves were based on a dilution series of total DNA extracted from a monoculture of *C. difficile* VPI 10463. The qPCR results were obtained using the CFX96 Touch Real-Time PCR Detections System (Bio-Rad Laboratories) and the reagent RealQ plus 2x Master Mix Green low Rox (Amplicon) as previously described [56].

### Pre-processing of fecal samples for separation of viruses and bacteria

Fecal samples from three different time points were included to investigate gut microbiome changes over time: baseline (before antibiotic treatment), before *C. difficile* infection (after antibiotic treatment), and at termination or at euthanization. This represented in total 142 fecal samples. Separation of the viruses and bacteria from the fecal samples generated a fecal pellet and fecal supernatant by centrifugation and 0.45 µm filtering as described previously [43], except the volume of fecal homogenate was adjusted to 5 mL using SM-buffer.

### Bacterial DNA extraction, sequencing and data pre-processing

The DNeasy PowerSoil Pro Kit (Qiagen) was used to extract bacterial DNA from the fecal pellet by following the instructions of the manufacturer. The final purified DNA was stored at -80 °C and the DNA concentration was determined using Qubit HS Assay Kit (Invitrogen) on the Qubit 4 Fluorometric Quantification device (Invitrogen). The bacterial community composition was determined by Illumina NextSeq-based high-throughput sequencing of the 16S rRNA gene V3-region, as previously described [43]. Quality control of reads, de-replicating, purging from chimeric reads and constructing Zero-radius Operational Taxonomic Units (zOTU) was conducted with the UNOISE pipeline [57] and taxonomy assigned with Sintax [58] using the EZtaxon for 16S rRNA gene database [59]. zOTU represents unique sequence variants where only sequence alignments with 100 % similarity are merged into the same zOTU. Code describing this pipeline can be accessed in https://github.com/jcame/Fastq_2_zOTUtable. The average sequencing depth after quality control (Accession: PRJEB58777, available at ENA) for the fecal 16S rRNA gene amplicons was 60,719 reads (min. 11,961 reads and max. 198,197 reads).

### Viral RNA/DNA extraction, sequencing and data pre-processing

The sterile filtered fecal supernatant was concentrated using Centrisart centrifugal filters with a filter cut-off at 100 kDA (Sartorius) by centrifugation at 1,500 x g at 4 °C (dx.doi.org/10.17504/protocols.io.b2qaqdse). The fecal supernatant (140 µL) was treated with 5 units of Pierce Universal Nuclease (ThermoFisher Scientific) for 10 minutes at room temperature prior to viral DNA extraction to remove free DNA/RNA molecules. The viral DNA/RNA was extracted from the fecal supernatants using the Viral RNA mini kit (Qiagen) as previously described [43,60]. Reverse transcription was executed with SuperScript IV VILO Master mix by following the instructions of the manufacturer and subsequently cleaned with DNeasy blood and tissue kit (Qiagen) by only following step 3-8 in the instructions from the manufacturer. In brief, the DNA/cDNA samples were mixed with ethanol, bound to the silica filter, washed two times, and eluted with 40 µL elution-buffer. Multiple displacement amplification (MDA, to include ssDNA viruses) using GenomiPhi V3 DNA amplification kit (Cytiva) and sequencing library preparation using the Nextera XT kit (Illumina) was performed at previously described [43], and sequenced using the Illumina NovaSeq platform at the sequencing facilities of Novogene (Cambridge, UK). The average sequencing depth of raw reads (Accession: PRJEB58777, available at ENA) for the fecal viral metagenome was 17,384,372 reads (min. 53,960 reads and max. 81,642,750 reads). Using Trimmomatic v0.35, raw reads were trimmed for adaptors and low quality sequences (<95 % quality, <50nt) were removed. High-quality reads were de-replicated and checked for the presence of PhiX control using BBMap (bbduk.sh) (https://www.osti.gov/servlets/purl/1241166). Virus-like particle-derived DNA sequences were subjected to within-sample *de novo* assembly-only using Spades v3.13.1 and contigs with a minimum length of 2,200 nt, were retained. Contigs from all samples were pooled and dereplicated by chimera-free species-level clustering at ∼95 % identity using the script described in [61], and available at https://github.com/shiraz-shah/VFCs. Contigs were classified as viral by VirSorter2 [62] (“full” categories | dsDNAphage, ssDNA, RNA, Lavidaviridae, NCLDV | viral quality = 1), VIBRANT [63] (High-quality | Medium-quality | Complete), CheckV [64] (High-quality | Medium-quality | Complete), and VirBot [65]. Any contigs not classified as viral by any of the 4 software’s were discarded. The taxonomical categories of “Other,” “Unclassified virus,” and “Unknown” that are used in the different figures are different entities. “Other” encompasses all remaining low abundance taxa not depicted in the plot. “Unknown” refers to contigs that may be viruses but lack specific data records confirming their viral origin, and “Unclassified virus” represents viruses that have been identified as having viral origin but could not be further classified. Taxonomy was inferred by blasting viral ORFs against a database of viral proteins created from the following: VOGDB v217 (vogdb.org), NCBI (downloaded 14/10/2023), COPSAC [61], and an RNA phage database [66], selecting the best hits with a minimum e-value of 10e^-6^. Phage-host predictions were done with IPhoP [67], which utilizes a combination of different host predictors. Following assembly, quality control, and annotations, reads from all samples were mapped against the viral (high-quality) contigs (vOTUs) using bowtie2 [68] and a contingency table of contig-length and sequencing-depth normalized reads, here defined as vOTU-table (viral contigs). Code describing this pipeline can be accessed in https://github.com/frejlarsen/vapline3. Mock phage communities (phage C2, T4, phiX174, MS2, and Phi6, Table S1) were used to both spike the FVT inoculums and as positive controls (normalized to ∼10^6^ PFU/mL for each phage) for virome sequencing to validate the sequencing protocol’s ability to include the different genome types of ssDNA, dsDNA, ssRNA, and dsRNA.

### Bioinformatics of bacterial and viral sequences and statistical analysis

The dataset was first cleaned to remove zOTU’s/viral contigs found in less than 5 % of the samples. Despite this, the resulting dataset retained over 99.8 % of the total reads. R version 4.3.2 was used for subsequent analysis and presentation of data. A minimum threshold of sequencing reads for the bacteriome and virome analysis was set to 2,200 reads and 15,000 reads, respectively. The main packages used were phyloseq [69], vegan [70], DESeq2 [71], ampvis2 [72], ggpubr, psych, igraph, ggraph, pheatmap, ComplexHeatmap, and ggplot2. Potential contaminations of viral contigs were removed by read count detected in negative controls through R package microDecon [73] (runs = 1, regressions = 1), and 35.1% of entries were removed. Cumulative sum scaling (CSS) was applied for the analysis of β-diversity. CSS normalization was performed using the R software using the metagenomeSeq package. α-diversity analysis (Shannon diversity-index) was based on raw read counts for bacteriome analysis, while the virome read counts were normalized on the basis of transcripts per million (TPM), and statistics were based on ANOVA. β-diversity was represented by Bray-Curtis dissimilarity and statistics were based on PERMANOVA. DESeq2 was used to identify differential abundant taxa on the summarized bacterial species level and viral contigs (vOTUs) level. The correlation heatmap between bacterial zOTUs and viral contigs (vOTUs) were calculated using pairwise Spearman’s correlations and FDR corrected. Cytokine levels, toxin levels, *C. difficile* abundance, and histology data were analyzed in R using linear models with saline as control group, while the log rank test was used to compare the survival distributions and FDR was used for corrections with multiple testing. Comparisons of means where used to calculate differences in PFU counts (https://www.medcalc.org/calc/comparison_of_means.php).

## RESULTS

We here hypothesized that different methodologies could be applied to overcome the challenges of donor variability and infection risks of eukaryotic viruses that are associated with FVT/FMT, while maintaining the treatment efficacy that previously have been reported for FVT/FMT treated recurrent *C. difficile* infection (rCDI) patients [5,12,13]. To produce “eukaryotic virus-free” fecal viromes, we developed methodologies that utilized fundamental differences in characteristics between eukaryotic viruses and phages: The majority of eukaryotic viruses are enveloped RNA viruses [28,29] and require eukaryotic hosts for replication, while the majority of phages are non-enveloped DNA viruses [28,30] and require bacterial hosts for replication. A solvent/detergent method was applied to inactivate enveloped viruses (FVT-SDT), pyronin-Y was used to inhibit replication of RNA viruses (FVT-PyT), and a chemostat propagated virome (FVT-ChP) was created to remove the majority of eukaryotic viruses by dilution [54]. These differently processed fecal viromes were as a proof-of-concept tested in a murine *C. difficile* infection model (Fig. 1) and compared with a saline solution (sham treatment), FMT, and untreated FVT (FVT-UnT). All treatments originated from the same intestinal donor content (and not from fecal pellets).

### Evaluation of methodologies applicability to inactivate enveloped and RNA viruses

The applicability of the solvent/detergent and pyronin-Y treatments to inactivate eukaryotic viruses while maintaining phage activity was tested by using phages representing different characteristics, such as enveloped (phi6) vs. non-enveloped (phiX174, T4, and C2) phage structure and ss/dsDNA (phiX174, T4, C2) vs. ss/dsRNA (MS2, phi6) genomes (Fig. 2A & 2B). The solvent/detergent treatment completely inactivated phage activity (as determined by plaque-forming units (PFU)/mL) of the enveloped phage phi6 from 10^9^ PFU/mL to below the detection limit (p < 0.0001). The activity of the non-enveloped phages phiX174 and T4 were largely unaffected with less than 0.1 log_10_ decrease, whereas, phage C2 showed a 1 log_10_ decrease in PFU/mL (p < 0.0001, Fig. 2A). Pyronin-Y was used to inactivate the replication of viruses harboring RNA genomes. Based on numerous combinations of pyronin-Y concentrations, temperatures, and incubation time, an overnight incubation at 40 °C with 100 µM pyronin-Y was chosen. This treatment reduced the ssRNA phage MS2 with 5 log_10_ PFU/mL (p < 0.0001) and dsRNA phage phi6 with more than 4 log_10_ PFU/mL (p < 0.0001) at 20 °C. Phi6 showed to be temperature sensitive since incubation at 40 °C alone inactivated (p < 0.0001) this enveloped phage. The plaque forming ability of phages C2 (dsDNA), T4 (dsDNA), and phiX174 (ssDNA) was unfortunately also affected by the pyronin-Y treatment at 40 °C, with a decrease of 1, 2.5, and 5 log_10_ PFU/mL (p < 0.0005, Fig. 2B), respectively. Thus, it would be expected that a notable fraction of the phages in the FVT will be inactivated using the pyronin-Y treatment. In a parallel study we showed that the chemostat propagation of fecal viromes led to a clear reduction in terms of relative abundance of eukaryotic viruses [54].

**Fig. 2:**
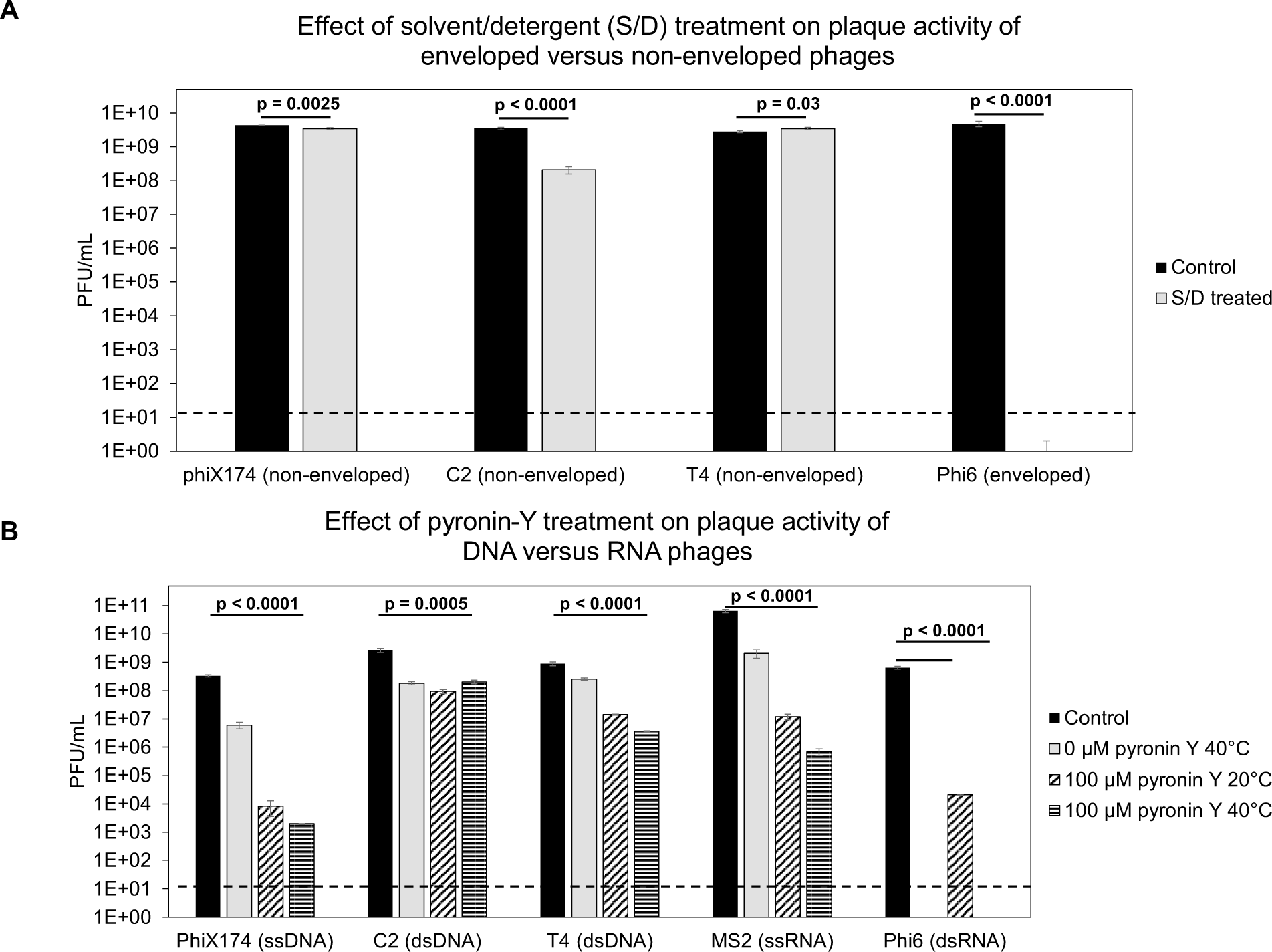
Evaluation of inactivation of phage activity (plaque-forming units, PFU/mL) with solvent/detergent or pyronin-Y treatment was evaluated on their respective bacterial hosts all performed in three replicates. Significance was evaluated with comparisons of means and t-test, with only considering the comparison between the control and the treatment applied in the study. A) Three non-enveloped phages (phiX174, C2, and T4) and one enveloped phage (phi6) were treated with solvent/detergent, and their plaque activity (plaque forming units per mL) was evaluated on their respective bacterial hosts. B) Phages representing ssDNA (phiX174), dsDNA (C2 and T4), ssRNA (MS2), and dsRNA (phi6) were treated with pyronin-Y, and their plaque activity (PFU/mL) at different incubation conditions was evaluated on their respective bacterial hosts. Dashed lines mark the detection limit of the applied assay.

### Fecal viromes maintained high treatment efficacy after inactivation of enveloped viruses

As a main endpoint parameter, the treatment efficacy of the FMT/FVTs, specifically preventing the mice from reaching the humane endpoint, was assessed in a murine *C. difficile* infection model (Fig. 1). The survival probability rate associated with the different treatments was evaluated using a Kaplan-Meier estimate (Fig. 3A) and compared to the mice treated with saline (2/7 mice). The analysis revealed a significantly improved survival rate (8/8 mice, *p* = 0.03) for mice treated with FVT-SDT, while the FVT-UnT (5/7 mice) and FVT-ChP (5/8 mice) treated mice showed tendencies (*p* = 0.24) of numerical, but non-significant, improvements of their survival rate. On the other hand, the FVT-PyT treated mice and, unexpectedly, the FMT treatment showed no improvement in survival rate (1/8 and 3/8 mice, respectively).

**Fig. 3:**
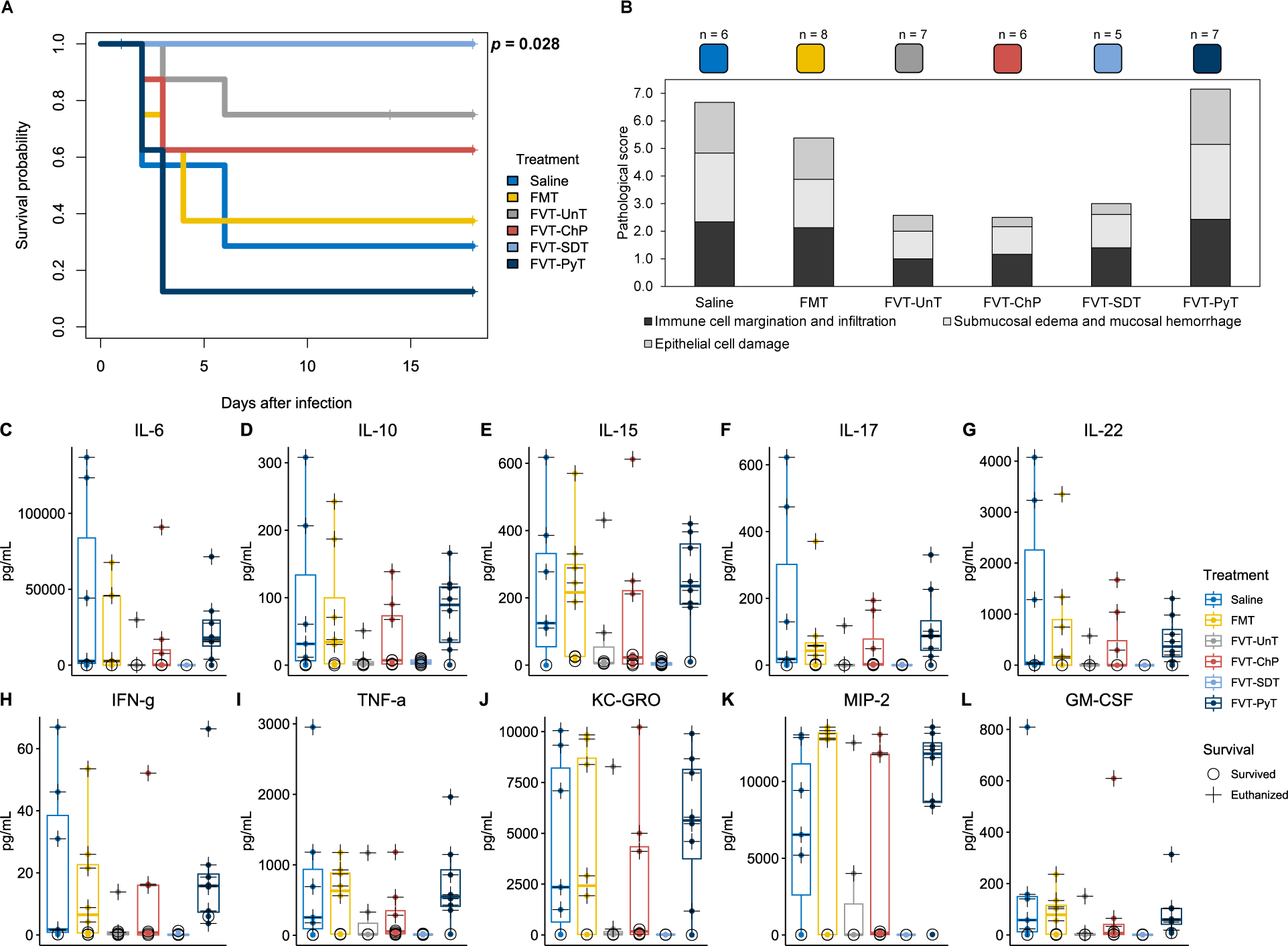
Overview of mouse phenotypic characteristics. A) Kaplan-Meier curve showing the survival probability of the mice that was associated to the different treatments when compared with the saline treated group. Pairwise comparisons between treatment groups with corrections for multiple testing were performed. B) Pathological score of cecum tissue evaluating the effect of the treatments’ ability to prevent *C. difficile* associated damage of the cecum tissue. C) to L) Showing the overall cytokine profile in the mouse cecum tissue of the different treatments. The euthanized (cross) mice are differentiated from the mice that survived (circle) the *C. difficile* infection and therefore represent two different time points. It was not possible to perform statistical analysis for the pathological score and cytokine profiles due to different timepoint of sampling of the euthanized and survived animals.

The pathological score (Fig. 3B, & Fig. S4A-J) and the levels of 10 pro- and anti-inflammatory cytokines of the cecum tissue were evaluated of both the mice that reached the humane endpoint and the mice that survived until study termination, which made it difficult to statistically evaluate these measures of the different sampling time points. The cecal histopathology and cytokine profiles therefore generally reflected whether the mice survived (with low or no inflammatory response when measured at study termination, marked with a circle) or were euthanized (with a high inflammatory response, marked with a cross) due to *C. difficile* infection (Fig. 3C-L). Thus, these measures were included to evaluate the animal’s disease status/recovery at termination or euthanization as well as the different FVT/FMT treatment abilities to increase the chances for the animals of not reaching the humane endpoint. However, a decrease in the average pathological score and cytokine levels supported the qualitative health evaluations (Fig. S4B-G) and the treatment efficacy associated with the improved survival rate of FVT-UnT, -SDT, and - ChP treated mice compared to the saline control, FMT, and FVT-PyT (Fig. 3B & Fig. S4A). The average pathological score at 6.7 of the saline treatment was in line with the original published *C. difficile* infection mouse model that reported a pathological score at 7.0 for *C. difficile* infected mice, compared to mice not infected with *C. difficile* that showed a score at 1.3 [31]. Overall, the FVT-SDT appeared as the superior treatment to prevent severe infections of *C. difficile*, since all 8 out of 8 mice did not reach the humane endpoint.

### Successful treatments impede *C. difficile* colonization and subsequent disease development

*C. difficile* abundance in feces was quantified using qPCR (Fig. 4A-C) to evaluate the infectious load at different time points. No *C. difficile* was detected before inoculation with *C. difficile* (Fig. 4A). The FVT-SDT treated mice exhibited an average of 2 log_10_ lower *C. difficile* abundance (*p* = 0.001) (gene copies per gram feces) compared to the saline treated mice before the 2^nd^ treatment, and the FVT-UnT (*p* = 0.013) and FVT-ChP (*p* = 0.039) treatments resulted in a 1.5 log_10_ lower abundance. This suggested that these three treatments effectively impeded *C. difficile* colonization in the gut. In contrast, the FMT and FVT-PyT treated mice had similar *C. difficile* abundance as the saline treated group. A possible clearance of *C. difficile* colonization was observed at study termination, as 7/8 FVT-SDT treated mice tested negative for *C. difficile*, while all other treatments showed persistency of *C. difficile* (Fig. 4C). The levels of the *C. difficile* associated toxin A/B were measured using an ELISA-based assay, which showed similar patterns as the qPCR data (Fig. 4D-F). Just before the 2^nd^ treatment, the toxin A/B levels in the FVT-SDT treated mice were significantly lower than those in the saline group (*p* < 0.05), and only 2 mice in the FVT-SDT group exhibited detectable toxin A/B. In contrast, toxin A/B was detected in all mice in the other FMT/FVT treatments and control (Fig. 4E). At termination, toxin A/B could not be detected in any of the FVT-SDT treated mice but was detected in a fraction of mice in the other treatment groups (Fig. 4F). The decrease in the abundance of *C. difficile*, the causing pathogenic agent (Fig. 4A-C), along with the diminished levels of toxin A/B (Fig. 4D-F) observed in the FVT-UnT, -SDT, and -ChP groups in comparison to mice treated with saline, aligns with the corresponding higher survival rates (Fig. 3A). This suggests a supportive relationship between reduced pathogen presence and toxin levels with improved overall chances of survival.

**Fig. 4:**
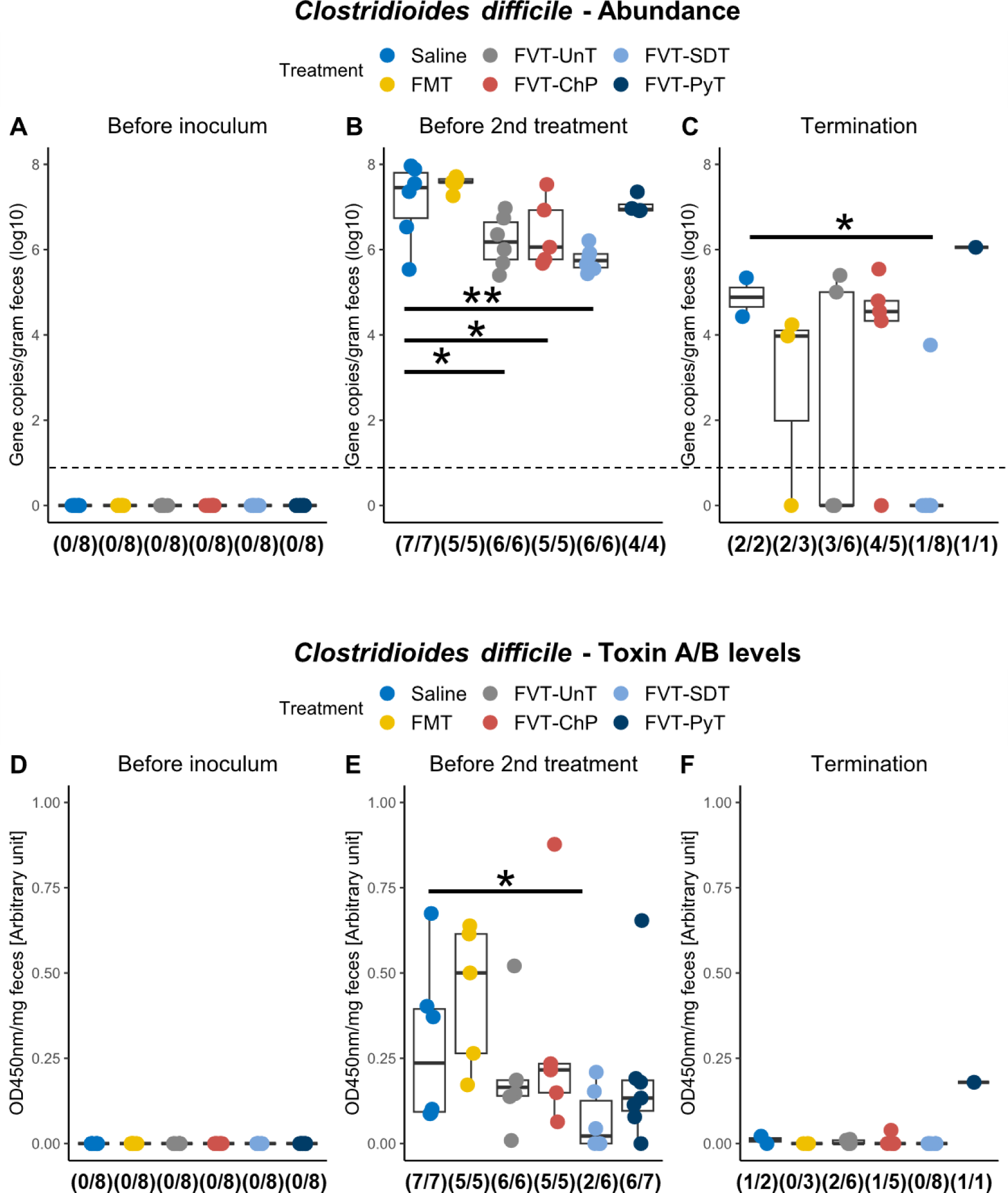
Evaluation of *C. difficile* abundance A) to C) by qPCR targeting the *toxA* gene and D) to F) the associated toxin A/B levels measured with ELISA on feces samples from three different time points: before *C. difficile* inoculum, before 2^nd^ treatment, and at study termination. The fraction below the boxplots highlights the number of mice that were detected positive of either *C. difficile* or toxin A/B. Dashed line marks the detection limit of the applied assay. *=*p*<0.05, **=*p*<0.01.

### Fecal virome treated with solvent/detergent supports recovery of the bacterial community in a dysbiotic gut microbiome

The gut microbiome analysis included three time points; baseline (before antibiotics), before *C. difficile* infection (after antibiotic treatment), and at planned termination or if the mice reached the humane endpoint prior study termination. The latter posed the inherent comparability challenge by the time difference between the euthanized mice and the mice that survived the infection (Fig. S5). Thus, these gut microbiome profiles reflected whether the mice survived the infection or were euthanized. Despite this, we deliberately included the gut microbiome data for all three time points to facilitate a comparative analysis for answering the following two main questions: 1) Did the different FVT/FMT treatments contribute to the restoration of the gut microbiome relative to baseline? 2) Which significant changes characterized the gut microbiome at the time of the euthanized and survived mice compared with the time point before the *C. difficile* infection? To do so, we verified that there were no initial differences (*p > 0.3)* in the bacterial and viral gut microbiome profiles at both baseline (before antibiotics) and before *C. difficile* infection (after antibiotic treatment) between the mice that were later either euthanized or survived the *C. difficile* infection (Fig. 5 & Fig. S6). The antibiotic intake through the drinking water was similar across the cages (Table S4). The overall bacterial composition (Bray-Curtis dissimilarity) and diversity (Shannon diversity index) were significantly different (*p* < 0.05) between the different time points (Fig. 5A-B), however the mice that survived the infection tended to be more similar to baseline, compared with the time before *C. difficile* infection and the euthanized mice. The bacterial taxonomic profile of the mice that had survived the infection was dominated by lactobacilli, *Prevotella, Clostridium sensu stricto, Bacteroides*, *Lachnospiraceae*, *Bifidobacterium*, *Akkermansia*, *Porphyromonadaceaea*, *Desulfovibrio, Parabacteroides,* and *Turicibacter*, which were also the dominant taxa in the mice at baseline (Fig. 5C-D), suggesting partial restoration of the gut microbiome profile in mice that survived the *C. difficile* infection. The bacterial taxonomic profile of the mice that were euthanized due to reaching the humane endpoint was consistently dominated by *Escherichia/Shigella*, *Enterococcus*, *Clostridioides*, *Bacteroides*, *Parasutterella,* and *Parabacteroides* (Fig. 5C-D). Except for the genus *Clostridioides*, these taxa were also among the more abundant before *C. difficile* infection, which indicated that the treatments at this time point had not restored the gut microbiome sufficiently after the antibiotic treatment. Two mice treated with FVT-PyT were colonized with 5-30% relative abundance of *Salmonella* spp. (Fig. S7), which may have contributed to increased disease severity leading to euthanasia of these two mice. The potential bacterial engraftment from the FMT inoculum to the FMT treated mice were analyzed at 16S rRNA gene amplicon level. The FMT treated mice that survived the infection were associated with a relative abundance of approx. 65 % that was also found in the FMT inoculum, compared with a relative abundance of approx. 15 % in the euthanized mice (Fig. S8A). This potential bacterial engraftment from the FMT inoculum was amongst others represented by *Clostridium sensu stricto* (Fig. 5C-D).

**Fig. 5:**
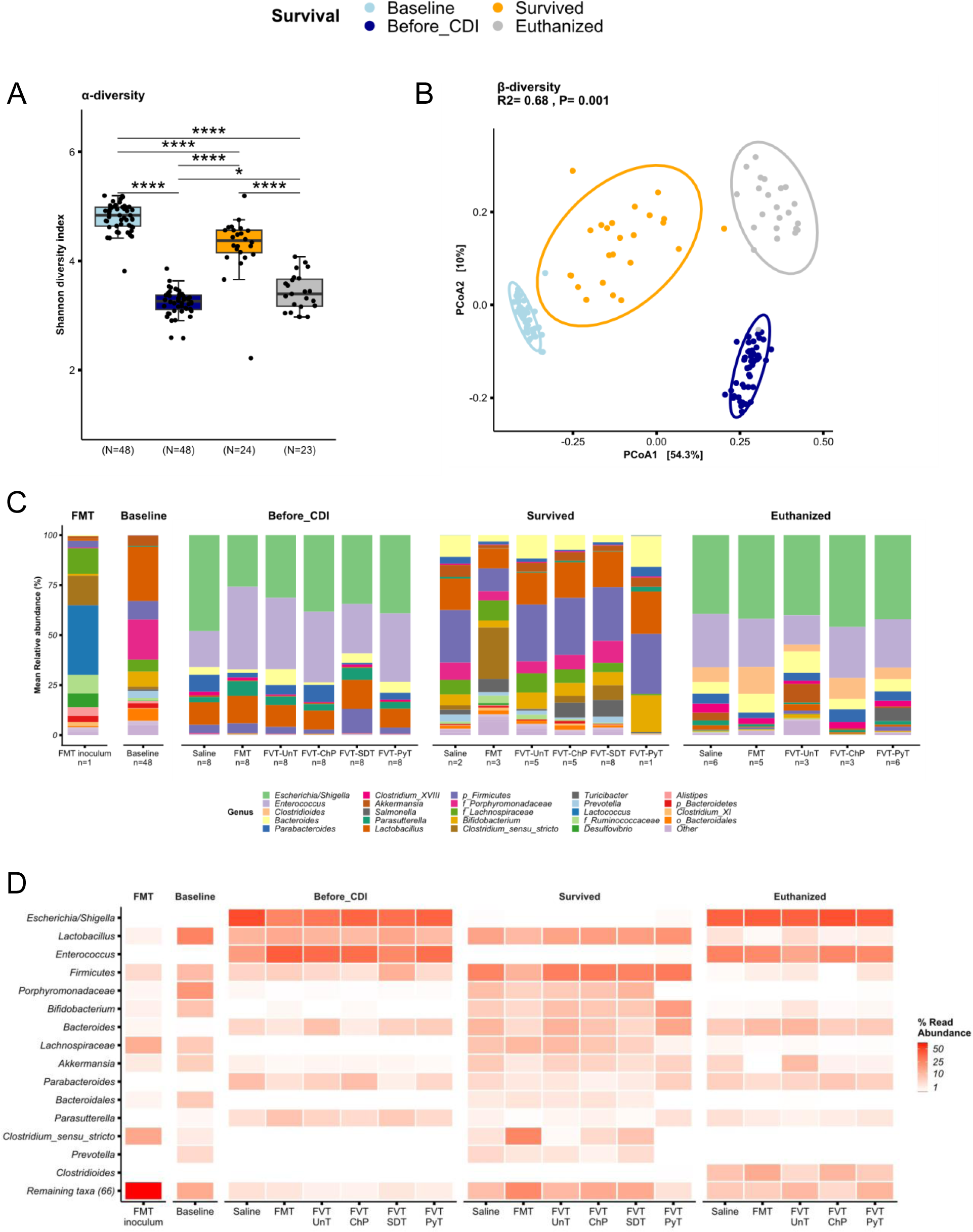
Bacteriome analysis based on 16S rRNA gene amplicon sequencing. A) The bacterial Shannon diversity-index (α-diversity) and B) Bray-Curtis dissimilarity based PCoA plot (β-diversity) at baseline (before antibiotic treatment), before *C. difficile* infection (after antibiotic treatment), for the mice that survived *C. difficile* infection (regardless of the treatment), and the euthanized mice. C) Bar plot and D) heatmap illustrating the bacterial relative abundance in percentage of the dominating bacterial taxa that was associated to the different mice that either survived the infection or were euthanized. The “n” below the bar plots highlights the number of mice of which the taxonomical average was based on. ***=p<0.005. ****=p<0.0005.

The viral composition and diversity of the mice that survived the infection were significantly different (p < 0.05) compared with the baseline, before C. difficile infection (Fig. 6A-B). The dominant viral taxa in all groups at all time points represented *Morgan*-, *Astro-, Alpa-, Mads-, Alma-, and Inesviridae*, while more than 60 % of the viral relative abundance could not be taxonomically assigned (Fig. 6C-E). The viral engraftment from the FVTs were also investigated at the viral metagenome level (viral contigs). Except for the FVT-UnT, the FMT/FVT treated mice that survived the infection were engrafted with a higher relative abundance of viral contigs that were also found in the different FMT/FVT inoculums, compared with the euthanized mice (Fig. S8B). This observation further indicated that the transfer of phages is likely associated with the treatment efficacy.

**Fig. 6:**
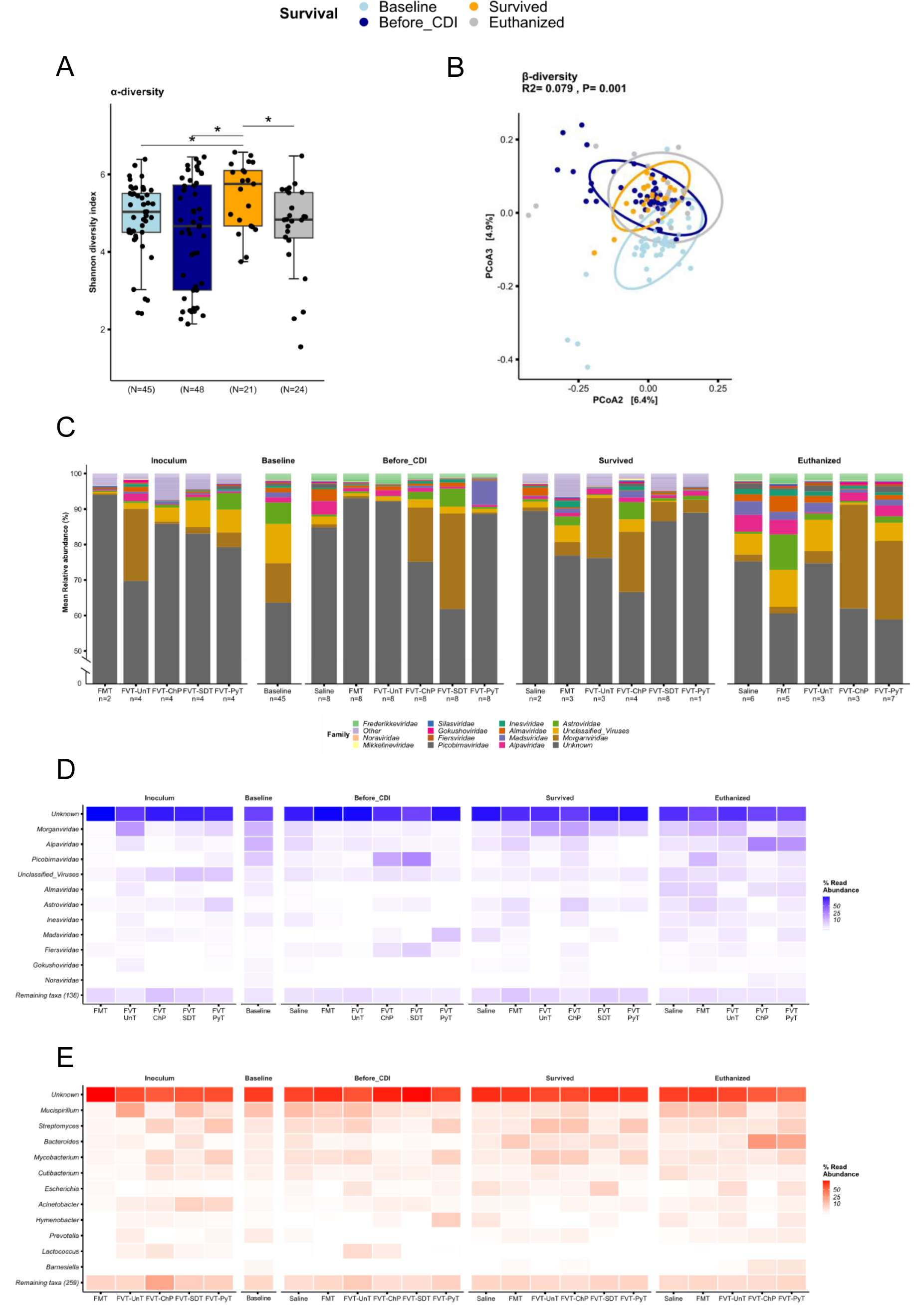
Metavirome analysis based on whole-genome sequencing. A) The viral Shannon diversity-index (α-diversity) and B) Bray-Curtis dissimilarity based PCoA plot (β-diversity) at baseline (before antibiotic treatment), before *C. difficile* infection (after antibiotic treatment), for the mice that survived *C. difficile* infection (regardless of the treatment), and the euthanized mice. C) Bar plot showing the relative abundance in percentage of the viral taxonomy (normalized on the basis of transcripts per million, TPM), and D) heatmap illustrating the relative abundance in percentage of the bacterial hosts that are predicted on the basis of the viral sequences. The “n” below the bar plots highlights the number of mice of which the taxonomical average was based on. **=p<0.01, ***=p<0.005.

Differential abundance testing was used to characterize the most significant gut microbiome changes of both the bacterial and viral component (relative abundance > 1% and *p* < 0.05) from the time before *C. difficile* infection (after antibiotic treatment) until the mice were euthanized or survived until the study termination (Fig. 7A-C). The mice that survived had significantly increased (*p* < 0.05) their relative abundance of bacterial taxa belonging to *Turicibacter*, *Clostridium sensu stricto*, *Akkermansia*, and *Clostridioides,* and a decrease in *Parasutterella, Parabacteroides, Enterococcus, Escherichia,* and *Bacteroides thetaiotaomicron* relative to before they were infected with *C. difficile* (Fig. 7A). The increase in *Clostridioides* is likely due to the persistence of *C. difficile* in the mice which also was detected by the quantitative analysis of *C. difficile* (Fig. 4C). The euthanized mice had significantly increased (*p* < 0.05) their relative abundance of especially *Clostridioides* (15 log_2_ fold change)*, Akkermansia*, and *Bacteroides* and a decrease in *Turicibacter*, lactobacilli, and *Parasutterella*, relative to before they were infected with *C. difficile* (Fig. 7B).

**Fig. 7:**
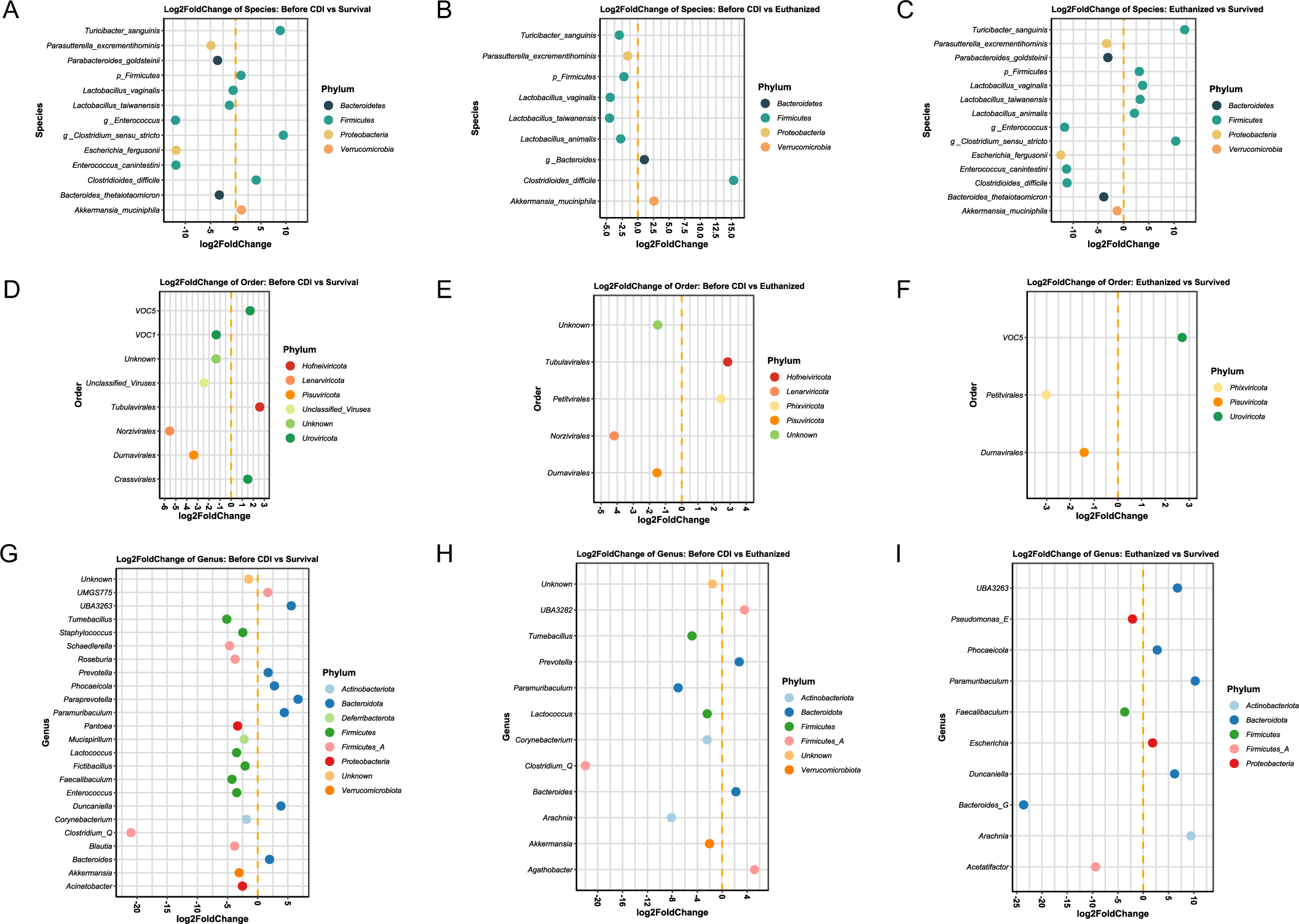
Differential abundance of significantly (*p* < 0.05) different A-C) bacterial taxa, D-F) viral taxa, and G-I) predicted bacterial hosts based on the viral sequences showing three comparisons: microbial taxa before *C. difficile* infection versus survived or euthanized mice, and the euthanized versus survived mice.

With regard to the phages (Fig. 7D-F), the surviving mice were characterized by an increase in viruses (*Vincent*-, *Sonia*-, *Rigmor*-, *Morgan*-, *Freja*-, *Ella*-, and *Christianviridae*) belonging to the viral order of *Crassvirales* and *Tubulavirales* relative to before they were infected with *C. difficile* (Fig. 7D). Whereas the euthanized mice mainly had increased their relative abundance of phages (*Nora*-, *Ines*-, *Gokusho*-, and *Alpaviridae*) belonging to the order *Petitvirales* (Fig. 7E). The viral metagenomes were used to predict potential bacterial hosts (Fig. 7G-I) with a recently developed machine learning framework that utilizes six different host prediction approaches [67]. Compared with the time point before *C. difficile* infection, the mice that survived the infection were characterized by a significant increase in (*p* < 0.05) of phages predicted to infect *Prevotella*, *Phocaeicola*, *Paraprevotella*, *Paramuribaculum*, *Duncaniella*, and *Bacteroides* members, and a decrease in vira predicted to infect *Tumebacillus*, *Staphyloccocus*, *Schaedlerella*, *Roseburia*, *Lactococcus*, *Fictibacillus*, *Faecalibacterium*, *Enterococcus*, *Clostridium*, *Blautia*, *Mucispririllum*, *Pantoea*, *Acinetobacter,* and *Akkermansia* (Fig. 7G). In contrast, the mice that were euthanized showed only an increase in phages predicted to infect *Prevotella*, *Bacteroides*, and *Agathobacter* and a decrease in *Tumebacillus*, *Lactococcus*, *Clostridium*, *Paramuribaculum, Coryinebacterium*, *Arachnia,* and *Akkermansia* (Fig. 7H). The overall host-phage relations between the relative abundance (> 0.1 %, *p* < 0.05) of bacterial zOTUs and viral contigs were assessed using Spearman’s correlations which showed clear clustering patterns (Fig. S9).

A bacterial and viral cluster A representing the taxa of *Enterococcus spp.*, *Salmonella*, *Clostridioides difficile*, *Escherichia fergusonii, Clostridium cocleatum, Bacteroides thetaiotamicron*, and *Parasutterella* positively correlated with mainly unknown viruses. Another bacterial and viral cluster B representing the taxa of *Prevotella,* lactobacilli*, Turicibacter, Clostridium spp., Porphyromonadaceaea, Lachnospiraceae, Bacteroides, Bifidobacterium,* and *Coriobacteriaceae* were positive correlated also with mainly unknown viruses. The bacterial genera in cluster B were associated with the mice that survived the infection, hence the unknown phages in this cluster may represent phages that had positively impacted the restoration of the GM compared with baseline. However, due to the limited viral classification it was not possible to detect clear evidence of specific host-phage relations that were driving the observed curative effects of FVT-UnT, -SDT, and -ChP.

Based on the above, it could be hypothesized that phages transferred along with the FVT-SDT treatment may have contributed to increased gut microbiome resilience against the *C. difficile* infection after the antibiotic treatment, and thereby impacting the mice’s ability to fight off the *C. difficile* infection. However, it remains uncertain whether the gut microbiome profile of the surviving mice has been similar in gut microbiome composition to that of the euthanized mice at a certain point of the *C. difficile* infection.

The eukaryotic viral profile was similar among the different FVT inoculates and constituted 0.1-3.0% of the total relative abundance and mainly represented RNA viruses (Fig. S10). However, it is important to note that the taxonomic resolution of eukaryotic viruses was insufficient to differentiate between treatments or outcomes in relation to the relative abundance of eukaryotic viruses (Fig. S10). It should also be emphasized that inactivation by dissolving the viral envelope or inhibiting replication of viruses in the FVT inoculums does not exclude the detection of viruses through sequencing and the metavirome analysis of FVT-SDT and –PyT can therefore not be used for validation whether specific viruses are inactivated or not.

## DISCUSSION

Here, we have developed methodologies to address the challenges of donor variability and risk of transferring pathogenic microorganisms when using FMT or FVT for treating gut-related diseases. A *C. difficile* infection mouse model was used as proof-of-concept. Inactivation of enveloped viruses through solvent/detergent treatment emerged as the superior method to modify fecal viromes while preserving treatment efficacy against *C. difficile* infection. The systemic inflammatory response observed during *C. difficile* infection is driven by *C. difficile* toxins that increase gut tissue permeability [74,75], making the gut more susceptible to other microbial infections [76]. Therefore, transferring untreated fecal donor viromes (including intact eukaryotic viruses) may result in additional inflammation due to microbes translocating through the damaged intestinal tissue. The majority of eukaryotic viruses are enveloped [28,29]. Hence, considering the promising prevention efficacy of *C. difficile* infection in the FVT-SDT group, treating fecal viromes with solvent/detergent prior to transfer to patients may be particularly relevant for diseases that are characterized by increased gut tissue permeability [76].

The lowest survival rate was observed with the RNA targeting compound pyronin-Y. During the initial evaluation of pyronin-Y’s ability to inactivate RNA phages, it became evident that DNA phages were also affected. The pyronin-Y treatment may have caused a reduction in phage activity, which could have impacted the efficacy of the treatment. This would align with several studies emphasizing the importance of high phage titers for successful treatment outcomes [77–80] and phages may play an important role in restoring gut microbiome balance following FMT or FVT [13,15–17,19,26,44,81].

FMT is linked to a treatment efficacy above 90 % in preclinical [32] and clinical *C. difficile* infection studies [5], however, the survival rate of FMT-treated mice was unexpectedly observed as similar to the saline group. The structure of the animal model does not allow us to assess whether this observation has biological relevance or represents a by-chance finding. Instead, we speculate in two potential explanations. First, the FMT inoculum contained approximately 20 % of *Clostridium sensu stricto* spp., which has been associated with *C. difficile*-positive calves [82] and to diarrhea in pigs [83]. The relatively high abundance of *Clostridium sensu stricto* spp. in the FMT inoculum may have counteracted the curative effects typically associated with FMT [13,32]. While the FMT and FVT inocula originated from the same donor material, the removal of bacteria during FVT processing may explain the higher survival rates observed with FVT-UnT, -ChP, and -SDT compared to FMT. This also suggests that even unsuitable fecal donor material for FMT could potentially be suitable for FVT, thus FVT-based treatments may have the potential to increase the probability in finding eligible donors, that have been reported as a challenging element of clinical FMT studies [7]. Secondly, a particularly controversial speculation could be that our FMT suspension was handled, sampled, and prepared anoxically inside an anaerobic chamber and stored/suspended in anoxic glycerol/PBS solutions. This is in strong contrast to the preparation of traditional FMT [84], which typically exposes the donor material to oxygen throughout the various preparation steps, from sampling to the final FMT product. It is well-acknowledged that the bacterial gut microbiome component mainly consists of strict and facultative anaerobes [85]. c of donor materials used for FMT unintentionally kills a vast number of oxygen-sensitive bacteria, reducing the load and diversity of viable bacterial cells. This, in turn, may decrease the chance of additional infections or other microbes translocating through damaged intestinal tissue that potentially could cause additional inflammation and tissue damage. In contrast, it would be expected that our anoxically handled FMT had a higher load and diversity of viable strict anaerobic bacteria [86,87], which may have counteracted the effect of the transferred enteric phages. However, it requires further studies to either confirm or reject this speculation.

Mice treated with FVT-UnT, -ChP, and -SDT showed a significant decrease in *C. difficile* abundance compared to those treated with saline, FMT, and FVT-PyT. We believe that phages transferred along with the FMT/FVT play a role in allowing commensal bacteria associated with a healthy state to compete with the infectious *C. difficile* strain, as well as commensal bacteria that can act as opportunistic pathogens. This was supported by phage engraftment from the FMT/FVT that was associated with mice surviving the infection and a cluster of unknown viruses that were positively correlated with bacteria reflecting a restored gut microbiome. However, the precise mechanisms underlying the gut microbiome modulating effects of FVT remain poorly understood. Nonetheless, several studies have established that the phage donor profile to some extent can be transmissible to the gut of patients suffering from *C. difficile* infections through FMT [13,44,81]. Our prior work demonstrated that FVT from lean mice could induce a shift in the gut microbiome composition of obese mice, resembling that of lean individuals [16].

Additionally, we recently illustrated how FVT originating from donors with a relatively high abundance of *A. muciniphila* could significantly elevate the relative abundance of the endogenous *A. muciniphila* in recipient mice [21], and another study showed that autochthonously transferred FVT protected against stress-associated behavior in mice [33]. These observations from independent research groups imply that the phenotypic traits of FVT donors may be transferred to recipients, possibly driven by the inclination of phages to establish ecosystems similar to their origins. This process may involve cascading events [26], as illustrated in a gnotobiotic mouse model where phage infections indirectly influenced the bacterial balance [88]. Consequently, the more complex viral community of FVT could similarly impact the bacterial ecosystem that influence the metabolome resulting in systemic changes, as seen in our previous study [16]. While the notion of such effects may seem counterintuitive given the commonly held belief in the strain-specific nature of phages, a recent study proposed that phages could interact with distantly related microbial hosts [89]. Phage satellites have also been suggested to contribute to broader host ranges [90,91]. Furthermore, the transfer of potentially beneficial metabolic genes from temperate phages to their bacterial hosts may enhance host competitiveness and contribute to overall microbiota changes [92,93]. These findings align with recent research demonstrating the significant influence of nutritional and host environments on the community ecology [94], suggesting that cascading events initiated by FVT could catalyze changes in the host environment. Beyond the bacteria-phage relations, the role of the immune system in gut health should not be underestimated. Recent evidence suggests that phages interact with the immune system through mechanisms like TLR3 and TLR9 [95,96] or other mechanisms resulting in the uptake by mammalian cells [97,98] A recent review has summarized the current understanding of phage immunogenicity, highlighting parallels with eukaryotic viruses [99,100]. Thus, stimulation of the immune system may represent another mechanism behind the effects of FVT.

While there were good indications that phages were a key component in the observed effects associated with the FVT-UnT, - SDT, and -ChP treated mice, it remains possible that metabolites or entities with a molecular size above 30 kDa (size cut-off of applied ultrafilter) may have contributed to the observed effects. These molecules could for instance be metabolites from lactobacilli [101], and *Akkermansia* spp. (pasteurized cell cultures) [102], bacteriocins with antimicrobial properties affecting the gut microbiome composition [103,104] or extracellular vehicles which have been shown to affect immune regulation during pregnancy [105,106], and may be involved in the etiology of inflammatory bowel diseases [107]. However, considering that most metabolites have a size less than 30 kDa [46,47], long-term colonization of donor phages in FMT studies [81,108,109], phages being associated to the treatment outcome of recurrent *C. difficile* infection [44,81], no reported effects of heat-treated FVT controls [15,110], and studies reporting beneficial effects of FVT in different etiology regimes [12,13,15,16,19] it suggests that the viral component of FVT constitute an important role. In addition, it could be speculated that the solvent-detergent treatment (FVT-SDT) likely have dissolved a certain fraction of lipid-based extracellular vesicles, and thereby further diminished their potential role in the treatment outcome using FVT-SDT.

The FVT preparation protocol does not remove microorganisms or other entities that can pass through the applied 0.45 µm filtration. Bacterial endospores exhibit a size range, with some as small as 0.25 µm, although their typical dimensions surpass 0.8 µm [111,112]. Similarly, certain bacterial species within the taxa of *Mycoplasma*, *Pelagibacter*, and *Actinobacteria* can attain sizes as small as 0.2 µm [113–118], while most bacteria generally range in length from 1 to 10 µm [119]. Hence, the use of a 0.45 µm sterile filtration is anticipated to eliminate the vast majority of bacterial cells and spores. This was supported by both the notably low colony-forming unit (CFU) counts observed in the FVTs under the investigated conditions (Table S2), and the 16S rRNA gene profile of the FVTs showing low read counts and/or no gut-associated bacteria in 3 out of 4 FVTs (Fig. S11). Choosing a smaller pore size like 0.22 µm can minimize contamination of bacteria but will cause in exclusion of large viruses and phages [120] and have been shown to negatively affect the abundance of common phages [121]. Thus, making 0.22 µm filtration an undesirable solution for FVT preparation. The abundance and significance of nano-sized bacteria in gut health remains sparsely investigated [122]. We can therefore neither confirm nor definitively rule out that these bacteria potentially have influenced the treatment outcomes of the FVTs.

The application of multiple displacement amplification (MDA) for 1.5 – 2.0 hours has been reported to compromise quantitative analysis of metagenomes by overestimating the abundance of ssDNA sequences [123,124], however, it has recently been shown that decreasing the time of whole genome amplification to 0.5 hours accommodate this bias to a level where it remain valid to compare inter-sample relative abundance of viruses [61]. The taxonomical classification of eukaryotic viruses mainly detected RNA viruses while only one DNA virus was detected. This would be in accordance with eukaryotic viruses being dominated by RNA viruses [28,29]. Phages are generally species or strain specific [14], but the limited bacterial taxonomical resolution, that are associated with 16S rRNA gene amplicon sequencing, restricts predicted host-phage correlations to the genus level of the bacteria.

A recent study showed that *C. difficile* senses the mucus layer since it moves towards the mucin glycan components due to chemotaxis, and that mucin-degrading bacteria like *Akkermansia muciniphila*, *Bacteroides thetaiotaomicron*, and *Ruminococcus torque* allow *C. difficile* to grow when co-cultured in culture media containing purified MUC2 but without glucose, despite *C. difficile* lacks the glycosyl hydrolases needed for degrading mucin glycans [125]. Interestingly, co-existence of these bacterial taxa may explain why the euthanized mice tended to lose most of the commensal bacteria like *Prevotella*, lactobacilli, *Turicibacter*, and *Bifidobacterium*, while *Akkermansia* and *Bacteroidetes* persisted. Preventive use of frequent administration of high doses of *A. muciniphila* has been shown to alleviate *C. difficile* infection associated symptoms in a similar mouse model [126], which together points in the direction of a co-existence rather than a symbiosis between *C. difficile* and mucin-degrading bacteria.

The high mortality rate associated with the included *C. difficile* VPI 10463 strain makes it valuable for assessing the main endpoint parameter of survival probability related to various FVTs. Euthanizing animals that reached the humane endpoint at different time points had naturally an impact on the statistical power and introduced challenges in evaluating time-dependent parameters such as cytokine profiles, histopathology, and comparable time series analysis of gut microbiome recovery. On the contrary, it is possible that if the surviving mice exhibited comparable histology, *C. difficile* abundance, cytokine profiles, toxin levels, and gut microbiome profiles they would have reached the humane endpoint at a similar time point as the euthanized mice. The animal model was designed as such to adhere to the 3Rs principles [39] (replacement, reduction, and refinement) by limiting the number of mice per group to 8 instead of employing several termination points for all treatment groups. In addition, the group size was evaluated as sufficient for screening the survival probability associated with the different FMT/FVT treatments. It would have provided additional insights of the role of phages in the FVT treatment of the *C. difficile* infection if UV and/or heat-treated FVT controls were included in the design of the animal model. However, due to the severity of the *C. difficile* infection model applied, it would not accommodate the principle of 3Rs (reduce) [39] to include additional animals considering that two previous studies have shown no effect of heat-treated FVT controls [15,110]. Thus, it would be extremely relevant to include such controls in future studies using animal models causing less severity in disease development.

As an alternative to phage-based therapies to restore a dysbiotic gut microbiome, a recent bacterial consortium (SER-109) has been approved by the FDA to treat rCDI [127,128], which highlights the potential of also using defined bacterial consortia in modulation of the gut microbiome. The treatment efficacy of SER-109 was found to be 28 % percentage points (88 %) increased compared with placebo (60 %) [127], while regular FMT in another study shows 57 % percentage points (90 %) enhanced treatment efficacy compared with placebo (33 %) [5]. This suggested that FMT may still be the preferred treatment strategy for rCDI depending on the patient group. Furthermore, the role on treatment outcome of induced prophages originating from the bacteria in the SER-109 consortium remains to be addressed. Additional studies are therefore necessary to be conducted for comparing phage-based treatments with FMT and bacterial consortia like SER-109.

A major challenge of *C. difficile* infection is the risk of recurrent infections [6]. It is therefore interesting to note that 7/8 FVT-SDT treated mice showed non-detectable of *C. difficile* at termination, indicating a decrease in the risk of recurrent infections when treated with a solvent/detergent modified fecal virome. The inherent challenges of variability and reproducibility in fecal donor material exist for both FMT and FVT [7,8]. Two independent studies have demonstrated how propagation of fecal inoculum in a chemostat-fermentation holds the potential for reproducing the enteric viral component [54,129]. Interestingly, the treatment efficacy and decrease in *C. difficile* infection-associated symptoms were also pronounced in mice treated with the chemostat propagated enteric virome. Therefore, it could be argued that the solvent/detergent methodology of fecal viromes, already approved by WHO as a safe procedure for treating blood plasma [50], holds the potential to complement FMT in the treatment of *C. difficile* infection in the short-term perspective. In the long-term perspective, a cost-effective, standardized, and reproducible chemostat propagated enteric phageome for *C. difficile* infection treatment may also have tremendous potential for phage-mediated treatment of other diseases associated with gut dysbiosis.

## CONCLUSION

The hypothesis of this proof-of-concept study was that different modifications of FVT had the potential to address the challenges of donor variability and infections risks that are associated with FVT/FMT. Especially two FVT modification strategies showed a significant effect in limiting the colonization of *C. difficile* in the infected mice and thereby increased their chance of survival. Inactivation of enveloped viruses through solvent/detergent treatment of the FVT appeared as a superior method to address the infection risks while preserving treatment efficacy against *C. difficile* infection. Also the chemostat propagated FVT showed promising potential as a methodology to address both donor variability and the infection risks, thus, overall confirming our initial hypothesis. Due to the natural limitations associated with the simplicity of the study, these results encourage additional preclinical studies to further validify the translatability and relevance of applying these concepts of FVT treatments in clinical settings.

## DECLARATIONS

### Animal ethical approval

All procedures involving handling of animals included in the *C. difficile* infection model (license ID: 2021-15-0201-00836) and donor animals (license ID: 2012-15-2934-00256) were approved and conducted in accordance with Directive 2010/63/EU and the Danish Animal Experimentation Act.

### Availability of data and material

All data associated with this study are present in the paper or the Supplementary Materials. All sequencing datasets are available in the ENA database under accession number PRJEB58777.

### Competing interests

All authors declare no financial or personal conflicts of interest.

### Funding

Funding was provided by: The Lundbeck Foundation with grant ID: R324-2019-1880 under the acronym “SafeVir”. The Novo Nordisk Foundation with grant ID: NNF-20OC0063874 under the acronym “PrePhage”.

### Author contributions

Conceptualization: TSR, DSN

Methodology: TSR, XM, SF, FL, SBL, KDT, AVM (veterinarian, supervising the health status monitoring), AB (scored histology images), JLCM, SA, KA, CHFH

Investigation: TSR, XM, SF, FL, AB, AVM, CHFH, AKH, DSN

Visualization: TSR, AB, XM

Funding acquisition: TSR, DSN

Project administration: TSR, DSN

Supervision: DSN, AKH

Writing – original draft: TSR

Writing – review & editing: All authors critically revised and approved the final version of the manuscript.

## Acknowledgement

We would like to express our gratitude to the staff of veterinarians and animal caretakers at the Department of Experimental Medicine (AEM, University of Copenhagen, Denmark) for their cooperation in housing, handling, and monitoring the mice. We would also like to extend our thanks to PhD Casper Normann Nurup for his assistance in the initial setup of the column chromatography, and to lab trainee Mariam Al-Batool Samir S Bagi for performing the ELISA assay. We would also like to acknowledge the Food & Health Open Innovation project (FOODHAY), granted by the Danish Ministry of Education and Research, for funding the qPCR equipment (Bio-Rad Laboratories CFX96) used in this study. Lastly, we are grateful for the funding bodies that financially have supported our work.

## Notes

### Competing Interest Statement

The authors have declared no competing interest.

### Summary of Updates

We have improved the metavirome analysis by a complete re-analysis of the viral sequencing data. Also, additional correlation and engraftment analysis has been performed to increase the resolution of the gut microbiome analysis. All figures have been updated, and several new supplemental figures have been added. We have added several new paragraphs to the discussion to expand the perspectives of the results as well as adressing the limitations of the study. Two additional co-authors (Frej Larsen and Xiaotian Mao) are added to the author list for their contributions. Frej have significantly improved the metavirome analysis by a complete re-analysis/assembly of the raw sequencing data. His effort has amongst other led to improved binning of viral contigs (vRhyme), viral taxonomic resolution (VirBot), and better host prediction using a recently published tool by Simon Roux (iPHoP). Xiaotian has contributed to the complete re-analysis, visualization, and interpretation of both the processed metavirome and 16S rRNA gene amplicon sequencing data.

## REFERENCES

1 Vijay A, Valdes AM. Role of the gut microbiome in chronic diseases: a narrative review.Eur J Clin Nutr. 2022;76:489–501.

2 Degruttola AK, Low D, Mizoguchi A, et al. Current understanding of dysbiosis in disease in human and animal models. Inflamm Bowel Dis. 2016;22:1137–50.

3 Vasilescu I-M, Chifiriuc M-C, Pircalabioru GG, et al. Gut dysbiosis and *Clostridioides difficile* infection in neonates and adults. Front Microbiol. 2022;12.

4 Dawkins JJ, Allegretti JR, Gibson TE, et al. Gut metabolites predict *Clostridioides difficile* recurrence. Microbiome. 2022;10.

5 Baunwall SMD, Andreasen SE, Hansen MM, et al. Faecal microbiota transplantation for first or second *Clostridioides difficile* infection (EarlyFMT): a randomised, double-blind, placebo-controlled trial. Lancet Gastroenterol Hepatol. 2022;7:1083–91.

6 Mullish BH, Quraishi MN, Segal JP, et al. The use of faecal microbiota transplant as treatment for recurrent or refractory *Clostridium difficile* infection and other potential indications: joint British Society of Gastroenterology (BSG) and Healthcare Infection Society (HIS) guidelines. Gut. 2018;67:1920–41.

7 Kassam Z, Dubois N, Ramakrishna B, et al. Donor screening for fecal microbiota transplantation. New England Journal of Medicine. 2019;381:2070–2.

8 Wilson BC, Vatanen T, Cutfield WS, et al. The super-donor phenomenon in fecal microbiota transplantation. Front Cell Infect Microbiol. 2019;9:1–11.

9 U.S. Food & Drug Administration. Important safety alert regarding use of fecal microbiota for transplantation and risk of serious adverse reactions due to transmission of multi-drug resistant organisms. 2019. https://www.fda.gov/vaccines-blood-biologics/safety-availability-biologics/important-safety-alert-regarding-use-fecal-microbiota-transplantation-and-risk-serious-adverse (accessed 13 June 2023)

10 U.S. Food & Drug Administration. Safety alert regarding use of fecal microbiota for transplantation and additional safety protections pertaining to monkeypox virus. 2022.

11 U.S. Food & Drug Administration. Safety alert regarding use of fecal microbiota for transplantation and risk of serious adverse events likely due to transmission of pathogenic organisms. 2020. https://www.fda.gov/vaccines-blood-biologics/safety-availability-biologics/safety-alert-regarding-use-fecal-microbiota-transplantation-and-risk-serious-adverse-events-likely (accessed 23 May 2023)

12 Kao DH, Roach B, Walter J, et al. Effect of lyophilized sterile fecal filtrate vs lyophilized donor stool on recurrent *Clostridium difficile* infection (rCDI): Prelimenary results from a randomized, double-blind pilot study. J Can Assoc Gastroenterol. 2019;2:101–2.

13 Ott SJ, Waetzig GH, Rehman A, et al. Efficacy of sterile fecal filtrate transfer for treating patients with *Clostridium difficile* infection. Gastroenterology. 2017;152:799–811.e7.

14 Cao Z, Sugimura N, Burgermeister E, et al. The gut virome: A new microbiome component in health and disease. EBioMedicine. 2022;81:104113.

15 Draper LA, Ryan FJ, Dalmasso M, et al. Autochthonous faecal viral transfer (FVT) impacts the murine microbiome after antibiotic perturbation. BMC Biol. 2020;18:173.

16 Rasmussen TS, Mentzel CMJ, Kot W, et al. Faecal virome transplantation decreases symptoms of type 2 diabetes and obesity in a murine model. Gut. 2020;69:2122–30.

17 Mao X, Larsen SB, Zachariassen LSF, et al. Transfer of modified fecal viromes alleviates symptoms of non-alcoholic liver disease and improve blood glucose regulation in an obesity mouse model. bioRxiv 2023. doi: 10.1101/2023.03.20.532903

18 Borin JM, Liu R, Wang Y, et al. Fecal virome transplantation is sufficient to alter fecal microbiota and drive lean and obese body phenotypes in mice. Gut Microbes. 2023;15.

19 Brunse A, Deng L, Pan X, et al. Fecal filtrate transplantation protects against necrotizing enterocolitis. ISME J. 2022;16:686–94.

20 Feng H, Xiong J, Liang S, et al. Fecal virus transplantation has more moderate effect than fecal microbiota transplantation on changing gut microbial structure in broiler chickens. Poult Sci. 2023;103282.

21 Rasmussen TS, Mentzel CMJ, Danielsen MR, et al. Fecal virome transfer improves proliferation of commensal gut *Akkermansia muciniphila* and unexpectedly enhances the fertility rate in laboratory mice. Gut Microbes. 2023;15.

22 Rasmussen TS, Jakobsen RR, Castro-Mejía JL, et al. Inter-vendor variance of enteric eukaryotic DNA viruses in specific pathogen free C57BL/6N mice. Res Vet Sci. 2021;136:1–5.

23 Lim ES, Zhou Y, Zhao G, et al. Early life dynamics of the human gut virome and bacterial microbiome in infants. Nat Med. 2015;21:1228–34.

24 Jansen D, Matthijnssens J. The emerging role of the gut virome in health and inflammatory bowel disease: Challenges, covariates and a viral imbalance. Viruses. 2023;15.

25 Doorbar J, Egawa N, Griffin H, et al. Human papillomavirus molecular biology and disease association. Rev Med Virol. 2015;25:2–23.

26 Rasmussen TS, Koefoed AK, Jakobsen RR, et al. Bacteriophage-mediated manipulation of the gut microbiome - promises and presents limitations. FEMS Microbiol Rev. 2020;44:507–21.

27 Zuppi M, Vatanen T, Wilson BC, et al. Phages modulate bacterial communities in the human gut following fecal microbiota transplantation. Res Sq 2024. doi: 10.21203/rs.3.rs-3883935/v1

28 Koonin E V., Dolja V V., Krupovic M. Origins and evolution of viruses of eukaryotes: The ultimate modularity. Virology. 2015;479–480:2–25.

29 Rey FA, Lok S-M. Common features of enveloped viruses and implications for immunogen design for next-generation vaccines. Cell. 2018;172:1319–34.

30 Sausset R, Petit MA, Gaboriau-Routhiau V, et al. New insights into intestinal phages. Mucosal Immunol. 2020;13:205–15.

31 Chen X, Katchar K, Goldsmith JD, et al. A mouse model of *Clostridium difficile*-associated disease. Gastroenterology. 2008;135:1984–92.

32 Seekatz AM, Theriot CM, Molloy CT, et al. Fecal microbiota transplantation eliminates *Clostridium difficile* in a murine model of relapsing disease. Infect Immun. 2015;83:3838– 46.

33 Ritz NL, Draper LA, Bastiaanssen TFS, et al. The gut virome is associated with stress-induced changes in behaviour and immune responses in mice. Nat Microbiol. 2024;9:359–76.

34 Larsen C, Andersen AB, Sato H, et al. Transplantation of fecal filtrate to neonatal pigs reduces post-weaning diarrhea: A pilot study. Front Vet Sci. 2023;10.

35 Offersen SM, Mao X, Spiegelhauer MR, et al. Fecal virome is sufficient to reduce necrotizing enterocolitis. ResearchSquare 2024. doi: 10.21203/rs.3.rs-3856457/v1

36 Percie du Sert N, Hurst V, Ahluwalia A, et al. The ARRIVE guidelines 2.0: Updated guidelines for reporting animal research. Br J Pharmacol. 2020;177:3617–24.

37 Gouliouris T, Brown NM, Aliyu SH. Prevention and treatment of *Clostridium difficile* infection. Clinical Medicine. 2011;11:75–9.

38 Perić A, Rančić N, Dragojević-Simić V, et al. Association between antibiotic use and hospital-onset *Clostridioides difficile* infection in University Tertiary Hospital in Serbia, 2011–2021: An ecological analysis. Antibiotics. 2022;11.

39 MacArthur Clark J. The 3Rs in research: a contemporary approach to replacement, reduction and refinement. Br J Nutr. 2018;120:S1–7.

40 Edwards AN, McBride SM. Isolating and purifying *Clostridium difficile* spores. Methods Mol Biol. 2016;1476:117–28.

41 Rasmussen TS, Streidl T, Hitch TCA, et al. *Sporofaciens musculi* gen. nov., sp. nov., a novel bacterium isolated from the caecum of an obese mouse. Int J Syst Evol Microbiol. 2019;71.

42 Darkoh C, DuPont HL, Norris SJ, et al. Toxin synthesis by *Clostridium difficile* is regulated through quorum signaling. mBio. 2015;6:e02569.

43 Rasmussen TS, de Vries L, Kot W, et al. Mouse vendor influence on the bacterial and viral gut composition exceeds the effect of diet. Viruses. 2019;11:435.

44 Zuo T, Wong SH, Lam K, et al. Bacteriophage transfer during faecal microbiota transplantation in *Clostridium difficile* infection is associated with treatment outcome. Gut. 2017;67:gutjnl-2017-313952.

45 Bibiloni R. Rodent models to study the relationships between mammals and their bacterial inhabitants. Gut Microbes. 2012;3:536–43.

46 Muthubharathi BC, Gowripriya T, Balamurugan K. Metabolomics: small molecules that matter more. Mol Omics. 2021;17:210–29.

47 Kim SJ, Kim SH, Kim JH, et al. Understanding metabolomics in biomedical research. Endocrinology and Metabolism. 2016;31:7–16.

48 Horowitz B, Bonomo R, Prince AM, et al. Solvent/detergent-treated plasma: a virus-inactivated substitute for fresh frozen plasma. Blood. 1992;79:826–31.

49 Remy MM, Alfter M, Chiem M-N, et al. Effective chemical virus inactivation of patient serum compatible with accurate serodiagnosis of infections. Clin Microbiol Infect. 2019;25:907.e7-907.e12.

50 WHO Health Product Policy and Standards Team. Guidelines on viral inactivation and removal procedures intended to assure the viral safety of human blood plasma products. 2004;WHO-TRS924-Annex4-1-82. https://www.who.int/publications/m/item/WHO-TRS924-Annex4 (accessed 2 January 2023)

51 Trescec A, Simić M, Branović K, et al. Removal of detergent and solvent from solvent-detergent-treated immunoglobulins. J Chromatogr A. 1999;852:87–91.

52 Darzynkiewicz Z, Kapuscinski J, Carter SP, et al. Cytostatic and cytotoxic properties of pyronin Y: relation to mitochondrial localization of the dye and its interaction with RNA. Cancer Res. 1986;46:5760–6.

53 Kapuscinski J, Darzynkiewicz Z. Interactions of pyronin Y(G) with nucleic acids. Cytometry. 1987;8:129–37.

54 Adamberg S, Rasmussen TS, Larsen SB, et al. Reproducible chemostat cultures to eliminate eukaryotic viruses from fecal transplant material. bioRxiv 2023. doi: 10.1101/2023.03.15.529189

55 Gillers S, Atkinson CD, Bartoo AC, et al. Microscale sample preparation for PCR of *C. difficile* infected stool. J Microbiol Methods. 2009;78:203–7.

56 Ellekilde M, Krych L, Hansen CHF, et al. Characterization of the gut microbiota in leptin deficient obese mice - Correlation to inflammatory and diabetic parameters. Res Vet Sci. 2014;96:241–50.

57 Edgar RC. Updating the 97% identity threshold for 16S ribosomal RNA OTUs. Bioinformatics. 2018;34:2371–5.

58 Edgar R. SINTAX: a simple non-Bayesian taxonomy classifier for 16S and ITS sequences. bioRxiv 2016. doi: 10.1101/074161

59 Kim O-S, Cho Y-J, Lee K, et al. Introducing EzTaxon-e: a prokaryotic 16S rRNA gene sequence database with phylotypes that represent uncultured species. Int J Syst Evol Microbiol. 2012;62:716–21.

60 Conceição-Neto N, Yinda KC, Van Ranst M, et al. NetoVIR: Modular approach to customize sample preparation procedures for viral metagenomics. *The Human Virome*. Methods in Molecular Biology. 2018:85–95.

61 Shah SA, Deng L, Thorsen J, et al. Expanding known viral diversity in the healthy infant gut. Nat Microbiol. 2023;8:986–98.

62 Guo J, Bolduc B, Zayed AA, et al. VirSorter2: a multi-classifier, expert-guided approach to detect diverse DNA and RNA viruses. Microbiome. 2021;9:37.

63 Kieft K, Zhou Z, Anantharaman K. VIBRANT: automated recovery, annotation and curation of microbial viruses, and evaluation of viral community function from genomic sequences. Microbiome. 2020;8:90.

64 Nayfach S, Camargo AP, Schulz F, et al. CheckV assesses the quality and completeness of metagenome-assembled viral genomes. Nat Biotechnol. 2021;39:578–85.

65 Chen G, Tang X, Shi M, et al. VirBot: an RNA viral contig detector for metagenomic data. Bioinformatics. 2023;39.

66 Neri U, Wolf YI, Roux S, et al. Expansion of the global RNA virome reveals diverse clades of bacteriophages. Cell. 2022;185:4023–4037.e18.

67 Roux S, Camargo AP, Coutinho FH, et al. iPHoP: An integrated machine learning framework to maximize host prediction for metagenome-derived viruses of archaea and bacteria. PLoS Biol. 2023;21:e3002083.

68 Langmead B, Salzberg SL. Fast gapped-read alignment with Bowtie 2. Nat Methods. 2012;9:357–9.

69 McMurdie PJ, Holmes S. phyloseq: an R package for reproducible interactive analysis and graphics of microbiome census data. PLoS One. 2013;8:e61217.

70 Dixon P. VEGAN, a package of R functions for community ecology. Journal of Vegetation Science. 2003;14:927–30.

71 Love MI, Huber W, Anders S. Moderated estimation of fold change and dispersion for RNA-seq data with DESeq2. Genome Biol. 2014;15:550.

72 Andersen KS, Kirkegaard RH, Karst SM, et al. ampvis2: an R package to analyse and visualise 16S rRNA amplicon data. bioRxiv 2018. doi: 10.1101/299537

73 McKnight DT, Huerlimann R, Bower DS, et al. microDecon: A highly accurate read-subtraction tool for the post-sequencing removal of contamination in metabarcoding studies. Environmental DNA. 2019;1:14–25.

74 Moore R, Pothoulakis C, LaMont JT, et al. *C. difficile* toxin A increases intestinal permeability and induces Cl-secretion. Am J Physiol. 1990;259:G165–72.

75 Rao K, Erb-Downward JR, Walk ST, et al. The systemic inflammatory response to *Clostridium difficile* infection. PLoS One. 2014;9:e92578.

76 Camilleri M. Leaky gut: mechanisms, measurement and clinical implications in humans. Gut. 2019;68:1516–26.

77 Sarker SA, Sultana S, Reuteler G, et al. Oral phage therapy of acute bacterial diarrhea with two coliphage preparations: a randomized trial in children from Bangladesh. EBioMedicine. 2016;4:124–37.

78 Jault P, Leclerc T, Jennes S, et al. Efficacy and tolerability of a cocktail of bacteriophages to treat burn wounds infected by *Pseudomonas aeruginosa* (PhagoBurn): a randomised, controlled, double-blind phase 1/2 trial. Lancet Infect Dis. 2019;19:35–45.

79 Duan Y, Llorente C, Lang S, et al. Bacteriophage targeting of gut bacterium attenuates alcoholic liver disease. Nature. 2019;575:505–11.

80 Nale JY, Spencer J, Hargreaves KR, et al. Bacteriophage combinations significantly reduce *Clostridium difficile* growth *in vitro* and proliferation *in vivo*. Antimicrob Agents Chemother. 2016;60:968–81.

81 Fujimoto K, Kimura Y, Allegretti JR, et al. Functional restoration of bacteriomes and viromes by fecal microbiota transplantation. Gastroenterology. 2021;160:2089–2102.e12.

82 Redding LE, Berry AS, Indugu N, et al. Gut microbiota features associated with *Clostridioides difficile* colonization in dairy calves. PLoS One. 2021;16:e0251999.

83 Zhu J-J, Gao M-X, Song X-J, et al. Changes in bacterial diversity and composition in the faeces and colon of weaned piglets after feeding fermented soybean meal. J Med Microbiol. 2018;67:1181–90.

84 Perez E, Lee CH, Petrof EO. A practical method for preparation of fecal microbiota transplantation. Methods in Molecular Biology. 2016;1476:259–67.

85 Hugon P, Dufour J-C, Colson P, et al. A comprehensive repertoire of prokaryotic species identified in human beings. Lancet Infect Dis. 2015;15:1211–9.

86 Bénard M V., Arretxe I, Wortelboer K, et al. Anaerobic feces processing for fecal microbiota transplantation improves viability of obligate anaerobes. Microorganisms. 2023;11:2238.

87 Papanicolas LE, Choo JM, Wang Y, et al. Bacterial viability in faecal transplants: Which bacteria survive? EBioMedicine. 2019;41:509–16.

88 Hsu BB, Gibson TE, Yeliseyev V, et al. Dynamic modulation of the gut microbiota and metabolome by bacteriophages in a mouse model. Cell Host Microbe. 2019;25:803–814.e5.

89 Hwang Y, Roux S, Coclet C, et al. Viruses interact with hosts that span distantly related microbial domains in dense hydrothermal mats. Nat Microbiol. 2023;8:946–57.

90 Barcia-Cruz R, Goudenège D, Moura de Sousa JA, et al. Phage inducible chromosomal minimalist island (PICMI), a family of satellites of marine virulent phages. bioRxiv 2023. doi: 10.1101/2023.07.18.549517

91 Eppley JM, Biller SJ, Luo E, et al. Marine viral particles reveal an expansive repertoire of phage-parasitizing mobile elements. Proceedings of the National Academy of Sciences. 2022;119.

92 Wittmers F, Needham DM, Hehenberger E, et al. Genomes from uncultivated Pelagiphages reveal multiple phylogenetic clades exhibiting extensive auxiliary metabolic genes and cross-family multigene transfers. mSystems. 2022;7.

93 Heyerhoff B, Engelen B, Bunse C. Auxiliary metabolic gene functions in pelagic and benthic viruses of the Baltic sea. Front Microbiol. 2022;13.

94 Weiss AS, Niedermeier LS, von Strempel A, et al. Nutritional and host environments determine community ecology and keystone species in a synthetic gut bacterial community. Nat Commun. 2023;14:4780.

95 Sweere JM, Van Belleghem JD, Ishak H, et al. Bacteriophage trigger antiviral immunity and prevent clearance of bacterial infection. Science (1979). 2019;363:eaat9691.

96 Gogokhia L, Buhrke K, Bell R, et al. Expansion of bacteriophages is linked to aggravated intestinal inflammation and colitis. Cell Host Microbe. 2019;25:285–299.e8.

97 Bichet MC, Chin WH, Richards W, et al. Bacteriophage uptake by mammalian cell layers represents a potential sink that may impact phage therapy. iScience. 2021;24:102287.

98 Bichet MC, Adderley J, Avellaneda-Franco L, et al. Mammalian cells internalize bacteriophages and use them as a resource to enhance cellular growth and survival. PLoS Biol. 2023;21:e3002341.

99 Champagne-Jorgensen K, Luong T, Darby T, et al. Immunogenicity of bacteriophages. Trends Microbiol. 2023;31:1058–71.

100 Kan L, Barr JJ. A mammalian cell’s guide on how to process a bacteriophage. Annu Rev Virol. 2023;10:183–98.

101 Teame T, Wang A, Xie M, et al. Paraprobiotics and postbiotics of probiotic lactobacilli, their positive effects on the host and action mechanisms: A review. Front Nutr. 2020;7.

102 Depommier C, Everard A, Druart C, et al. Supplementation with *Akkermansia muciniphila* in overweight and obese human volunteers: a proof-of-concept exploratory study. Nat Med. 2019;25:1096–103.

103 Dabour N, Zihler A, Kheadr E, et al. *In vivo* study on the effectiveness of pediocin PA-1 and *Pediococcus acidilactici* UL5 at inhibiting *Listeria monocytogenes*. Int J Food Microbiol. 2009;133:225–33.

104 Umu ÖCO, Bäuerl C, Oostindjer M, et al. The potential of class II bacteriocins to modify gut microbiota to improve host health. PLoS One. 2016;11:e0164036.

105 Abrahams VM, Straszewski-Chavez SL, Guller S, et al. First trimester trophoblast cells secrete Fas ligand which induces immune cell apoptosis. Mol Hum Reprod. 2004;10:55– 63.

106 Hedlund M, Stenqvist A-C, Nagaeva O, et al. Human placenta expresses and secretes NKG2D ligands via exosomes that down-modulate the cognate receptor expression: evidence for immunosuppressive function. The Journal of Immunology. 2009;183:340– 51.

107 Leonetti D, Reimund J-MM, Tesse A, et al. Circulating microparticles from Crohn’s disease patients cause endothelial and vascular dysfunctions. PLoS One. 2013;8:e73088.

108 Broecker F, Russo G, Klumpp J, et al. Stable core virome despite variable microbiome after fecal transfer. Gut Microbes. 2017;8:214–20.

109 Draper LA, Ryan FJ, Smith MK, et al. Long-term colonisation with donor bacteriophages following successful faecal microbial transplantation. Microbiome. 2018;6:1–9.

110 Reyes A, Wu M, McNulty NP, et al. Gnotobiotic mouse model of phage-bacterial host dynamics in the human gut. Proceedings of the National Academy of Sciences. 2013;110:20236–41.

111 Wolken WAM, Tramper J, van der Werf MJ. What can spores do for us? Trends Biotechnol. 2003;21:338–45.

112 Colas de la Noue A, Natali F, Fekraoui F, et al. The molecular dynamics of bacterial spore and the role of calcium dipicolinate in core properties at the sub-nanosecond time-scale. Sci Rep. 2020;10:8265.

113 Marshall WF, Young KD, Swaffer M, et al. What determines cell size? BMC Biol. 2012;10:101.

114 Young M, Artsatbanov V, Beller HR, et al. Genome sequence of the fleming strain of *Micrococcus luteus*, a simple free-living actinobacterium. J Bacteriol. 2010;192:841–60.

115 Ghuneim L-AJ, Jones DL, Golyshin PN, et al. Nano-sized and filterable bacteria and archaea: Biodiversity and function. Front Microbiol. 2018;9.

116 Hahn MW, Lünsdorf H, Wu Q, et al. Isolation of novel ultramicrobacteria classified as *Actinobacteria* from five freshwater habitats in Europe and Asia. Appl Environ Microbiol. 2003;69:1442–51.

117 Cazanave C, Manhart LE, Bébéar C. *Mycoplasma genitalium*, an emerging sexually transmitted pathogen. Med Mal Infect. 2012;42:381–92.

118 López-Pérez M, Haro-Moreno JM, Iranzo J, et al. Genomes of the ‘*Candidatus Actinomarinales*’ order: Highly streamlined marine epipelagic *Actinobacteria*. mSystems. 2020;5.

119 Volland J-M, Gonzalez-Rizzo S, Gros O, et al. A centimeter-long bacterium with DNA contained in metabolically active, membrane-bound organelles. Science (1979). 2022;376:1453–8.

120 Berg M, Roux S. Extreme dimensions - how big (or small) can tailed phages be? Nat Rev Microbiol. 2021;19:407.

121 Castro-Mejía JL, Muhammed MK, Kot W, et al. Optimizing protocols for extraction of bacteriophages prior to metagenomic analyses of phage communities in the human gut. Microbiome. 2015;3:64.

122 Binda C, Lopetuso LR, Rizzatti G, et al. *Actinobacteria*: A relevant minority for the maintenance of gut homeostasis. Digestive and Liver Disease. 2018;50:421–8.

123 Marine R, McCarren C, Vorrasane V, et al. Caught in the middle with multiple displacement amplification: the myth of pooling for avoiding multiple displacement amplification bias in a metagenome. Microbiome. 2014;2:3.

124 Yilmaz S, Allgaier M, Hugenholtz P. Multiple displacement amplification compromises quantitative analysis of metagenomes. Nat Methods. 2010;7:943–4.

125 Engevik MA, Engevik AC, Engevik KA, et al. Mucin-degrading microbes release monosaccharides that chemoattract *Clostridioides difficile* and facilitate colonization of the human intestinal mucus layer. ACS Infect Dis. 2021;7:1126–42.

126 Wu Z, Xu Q, Gu S, et al. *Akkermansia muciniphila* ameliorates *Clostridioides difficile* infection in mice by modulating the intestinal microbiome and metabolites. Front Microbiol. 2022;13:841920.

127 Feuerstadt P, Louie TJ, Lashner B, et al. SER-109, an oral microbiome therapy for recurrent *Clostridioides difficile* infection. New England Journal of Medicine. 2022;386:220–9.

128 Khanna S, Sims M, Louie TJ, et al. SER-109: An oral investigational microbiome therapeutic for patients with recurrent Clostridioides difficile infection (rCDI). Antibiotics. 2022;11:1234.

129 Santiago-Rodriguez TM, Ly M, Daigneault MC, et al. Chemostat culture systems support diverse bacteriophage communities from human feces. Microbiome. 2015;3:58.

