## Supplemental figures for "Overcoming donor variability and risks associated with fecal microbiota transplants through bacteriophage-mediated treatments"

### Supplementary tables

Table S1: List of included bacterial and phage strains along with relevant strain-specific information and growth conditions that was used for plaque assays. The same phages were also used as a mock community (positive control) and spiking for the metavirome sequencing where the phages where normalized to 10^6^ PFU/mL for each phage.

**A**

| **Phage** | **DSM ID** | **Family** | **Genome type** | **Bacterial host** | **Strain ID** | **DSM ID** | **Incubation temp. (°C)** | **Media** |
| --- | --- | --- | --- | --- | --- | --- | --- | --- |
| C2 | In house | *Ceduovirus* c2 | dsDNA | *Lactococcus lactis* | MG1363 | DSM 4366 | 30 | M17 |
| T4 | 4505 | *Straboviridae* | dsDNA | *Escherichia coli* | Luria | DSM 613 | 37 | LB |
| PhiX174 | 4497 | *Microviridae* | ssDNA | *Escherichia coli* | PC 0886 | DSM 13127 | 37 | BHI |
| Phi6 | 21518 | *Cystoviridae* | dsRNA | *Pseudomonas* sp. | HER 1102 | DSM 21482 | 25 | TSB |
| MS2 | 13767 | *Fiersviridae* | ssRNA | *Escherichia coli* | W1485 | DSM 5695 | 37 | NZCYM + 2mg/L streptomycin |

Table S2: The FVTs used in the current study were anaerobically incubated on non-selective Gifu anaerobic medium (GAM) for 14 days at 37 °C. Here we observed the below colony forming units (CFU) counts for each of the applied FVTs.

|  | **CFU Count** | | **CFU/mL** |
| --- | --- | --- | --- |
| **Sample/Replicate** | Replicate 1 | Replicate 2 | **Avg.** |
| FVT-SDT | 1 | 1 | 2.5 x 10^1^ |
| FVT-UnT | 0 | 0 | 0 or ≤ 2.5 x 10^1^ |
| FVT-ChP | 32 | 25 | 7.1 x 10^2^ |
| FVT-PyT | 1 | 1 | 2.5 x 10^1^ |
| SM-buffer (Saline) | 0 | 0 | 0 or ≤ 2.5 x 10^1^ |

Table S3: List of components and their associated concentration (g/L) in the growth medium used for chemostat propagation of the chemostat propagated fecal virome (FVT-ChP).

| **Class description** | **Ingredients** | **g/L** |
| --- | --- | --- |
| Protein | Tryptone | 3 |
| Carbohydrate | Starch, Corn | 5 |
| Mineral | Sodium Chloride | 0.527 |
| Mineral | Magnesium Sulfate, Heptahydrate | 0.036 |
| Mineral | Ammonium Molybdate Tetrahydrate | 0.001 |
| Mineral | MgCl2 | 0.002 |
| Mineral | FeSO4*7H2O | 0.0001 |
| Mineral | CaCl2 | 0.009 |
| Mineral | MnSO4*H2O | 0.003 |
| Mineral | ZnSO4*7H2O | 0.001 |
| Mineral | CoSO4*7H2O | 0.001 |
| Mineral | CuSO4*5H2O | 0.001 |
| Vitamin | Nicotinamide | 0.00025 |
| Vitamin | Biotin | 0.00025 |
| Vitamin | Pantothenic Acid, d, Calcium (a.k.a. B5) | 0.00025 |
| Vitamin | Pyridoxine HCl (a.k.a. B6) | 0.0005 |
| Vitamin | Riboflavin (a.k.a. B2) | 0.00025 |
| Vitamin | Thiamine HCl (a.k.a. B1) | 0.00025 |
| Vitamin | Folic Acid | 0.00025 |
| Vitamin | Menadione Sodium Bisulfite | 0.0005 |
| Vitamin | Pyridoxal-HCl | 0.0002 |
| Fiber | Apple pectin | 2 |
| Fiber | xylan (corn core) | 2 |
| Fiber | Arabinogalactan (larch wood) | 2 |
| Fiber | inulin HP (chikory) | 1 |
| Peptides/nucleic acids/vitamins | Yeast extract | 3 |
| Mucin | Mucin (porcine gastric type II) | 4 |
| Buffer | K2HPO4, pH 6.4 | 2.4 |
| Buffer | KH2PO4, pH 6.4 | 4.9 |
| Lipids | Tween-80, ml | 0.5 |
| Bile salt | Bile salts | 0.5 |
| Buffer/carbonate | NaHCO3 | 2 |
| amino acid | Cys-HCl fresh | 0.5 |
| Hemin | Hemin | 0.005 |
| Redox | Na-thioglycolate | 0.5 |
| Organic acid | Acetic acid | 0.3 |

Table S4: List of antibiotic (AB) water consumption. The volumes were calculated as the average of each cage.

| **Experimental group** | **Cage no.** | **Average AB water consumption per mouse (mL)** |
| --- | --- | --- |
| Saline | 1 | 8.0 |
| FMT | 2 | 11.8 |
| FVT-UnT | 3 | 8.0 |
| FVT-ChP | 4 | 12.0 |
| FVT-SDT | 5 | 10.8 |
| FVT-PyT | 6 | 10.0 |
| Saline | 7 | 6.0 |
| FMT | 8 | 7.8 |
| FVT-UnT | 9 | 9.8 |
| FVT-ChP | 10 | 10.5 |
| FVT-SDT | 11 | 11.3 |
| FVT-PyT | 12 | 11.3 |
