## Supplemental tables for "Overcoming donor variability and risks associated with fecal microbiota transplants through bacteriophage-mediated treatments"

**Supplementary figures**

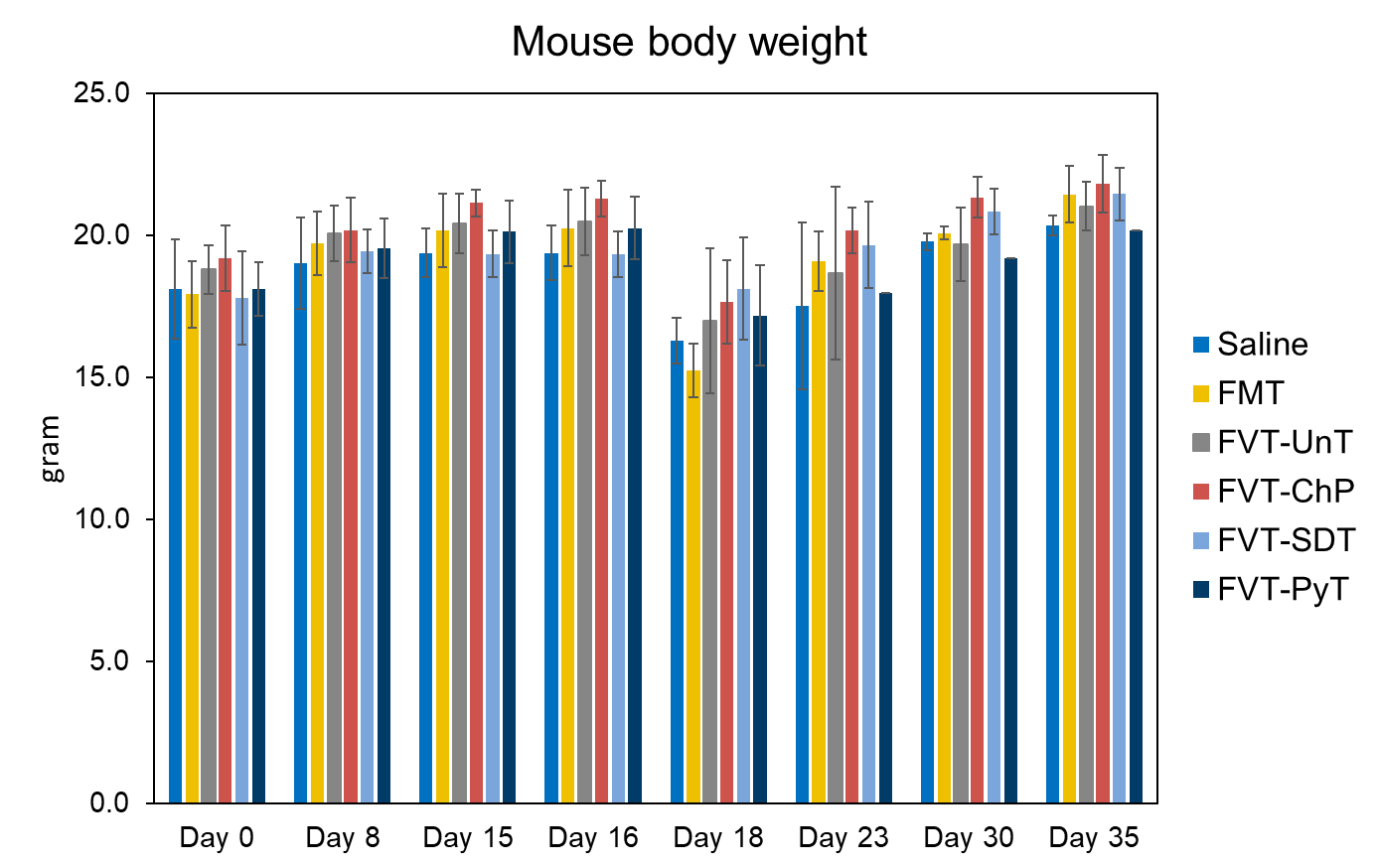

Fig. S1: Mouse body weight measured at arrival (day 0), just before antibiotic treatment (day 8), just before *C. difficile* infection (day 15), just before 1^st^ treatment (day 16), just before 2^nd^ treatment (day 18), one week after 2^nd^ treatment (day 23), two weeks after 2^nd^ treatment (day 30), and termination (day 35).

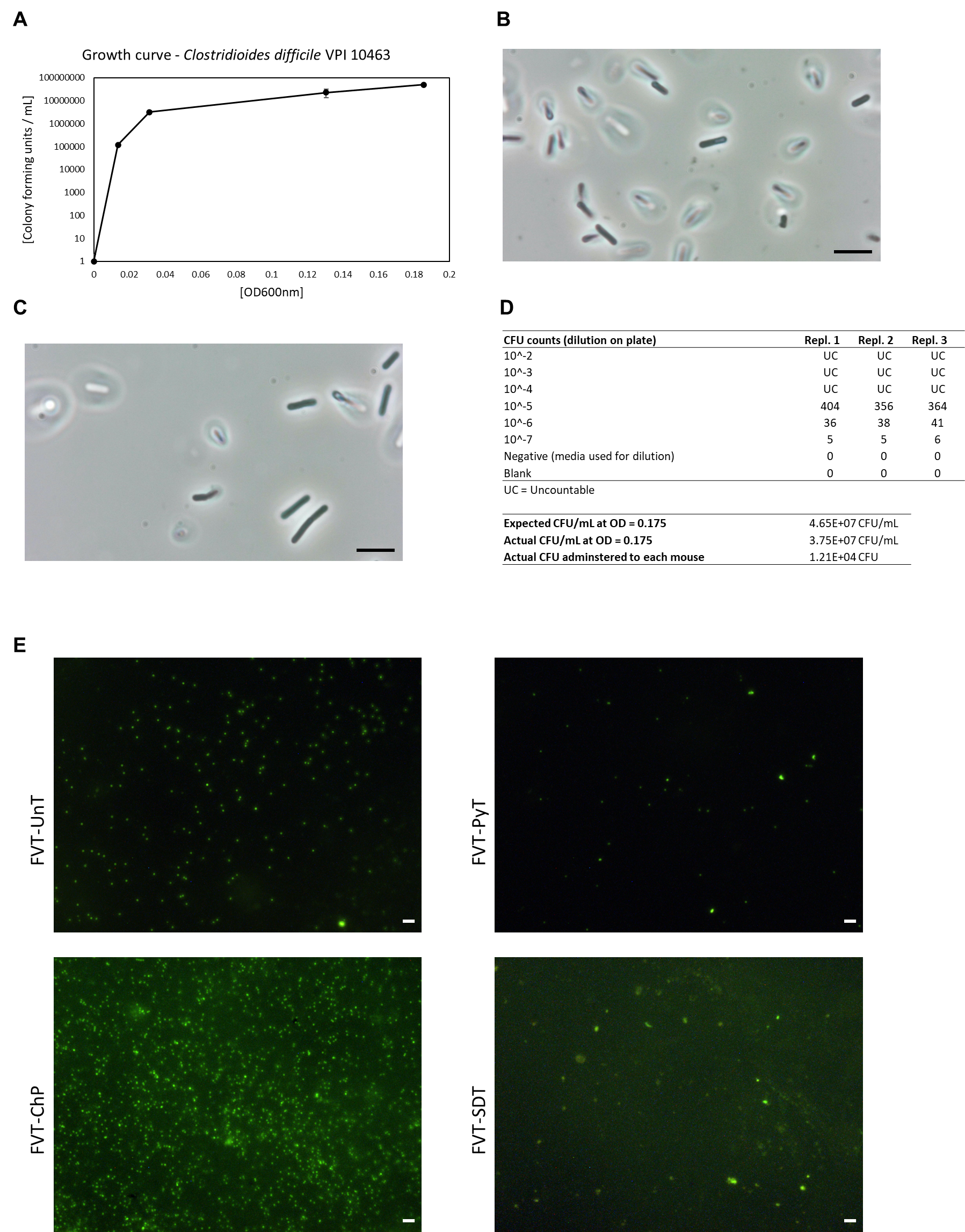

Fig. S2: A) Bacterial calibration curve showing the correlation between OD_600nm_ and colony forming unite per mL (CFU/mL) of *C. difficile* VPI 10463 and B) and C) phase contrast microscopy confirming the expected cell morphology of *C. difficile* VPI 10463. D) CFU counts in technical replicates of the applied *C. difficile* inoculum and the total CFU that was transferred to each mouse. E) Epifluorescence microscopy images of the different applied FVT viromes, before being normalized to similar virus-like particles(VLP)/mL concentrations that were stained with SYBR Gold to count the VLP/mL. Images were taken using a 100x magnification objective. Scale bar for the bacteria is 4 µm in C) and D) and for VLPs in E) the scale bar is 1 µm.

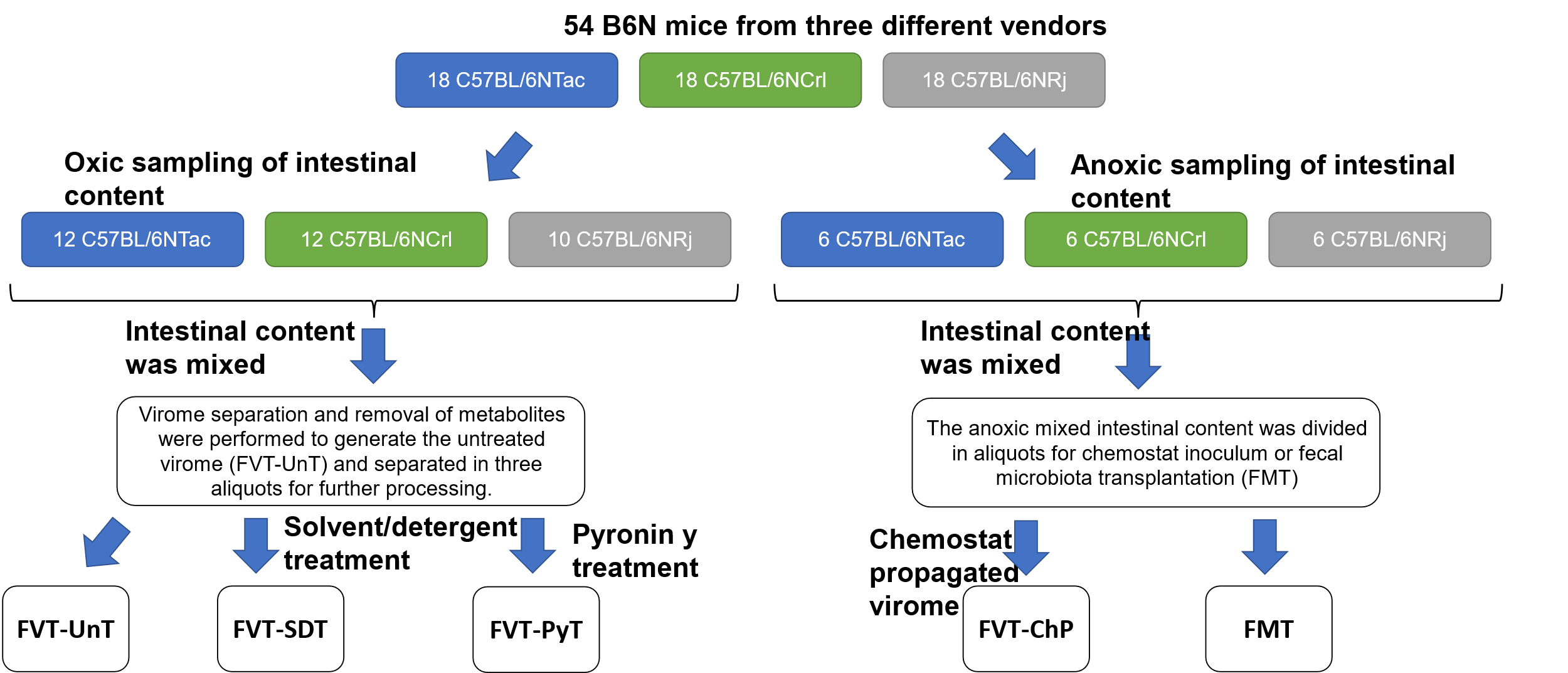

Fig. S3: Flow diagram illustrating the origin of the intestinal donor content and the processing steps to generate the FVTs with different methodologies. Two C57BL/6NRj mice were euthanized due to malocclusion-associated malnutrition. Then 52 mice from 3 different vendors were sacrificed and their intestinal content from cecum and colon was collected, however, 34 mice were sampled in oxic conditions, while 18 mice were sampled in anoxic conditions to maintain the viability of the strict anaerobic bacterial gut microbiome members. The atmospheric conditions were maintained throughout the process. Regardless of vendor, the intestinal content was mixed. Virome separation was performed for the oxic handled fecal mixture and the extracted fecal viromes were divided in three aliquots that represented 1) the untreated fecal virome (FVT-UnT), and 2) the further processing with solvent/detergent treatment (FVT-SDT) or 3) pyronin-Y treatment (FVT-PyT). Anoxic handled fecal mixture was divided in two aliquots for use as fecal microbiota transplantation (FMT) in the CDI mouse model, or as inoculum for fecal virome propagation in a chemostat setup (FVT-ChP). Also, the FVT-ChP underwent virome separation to remove most metabolites as well as bacteria and other larger microbes.

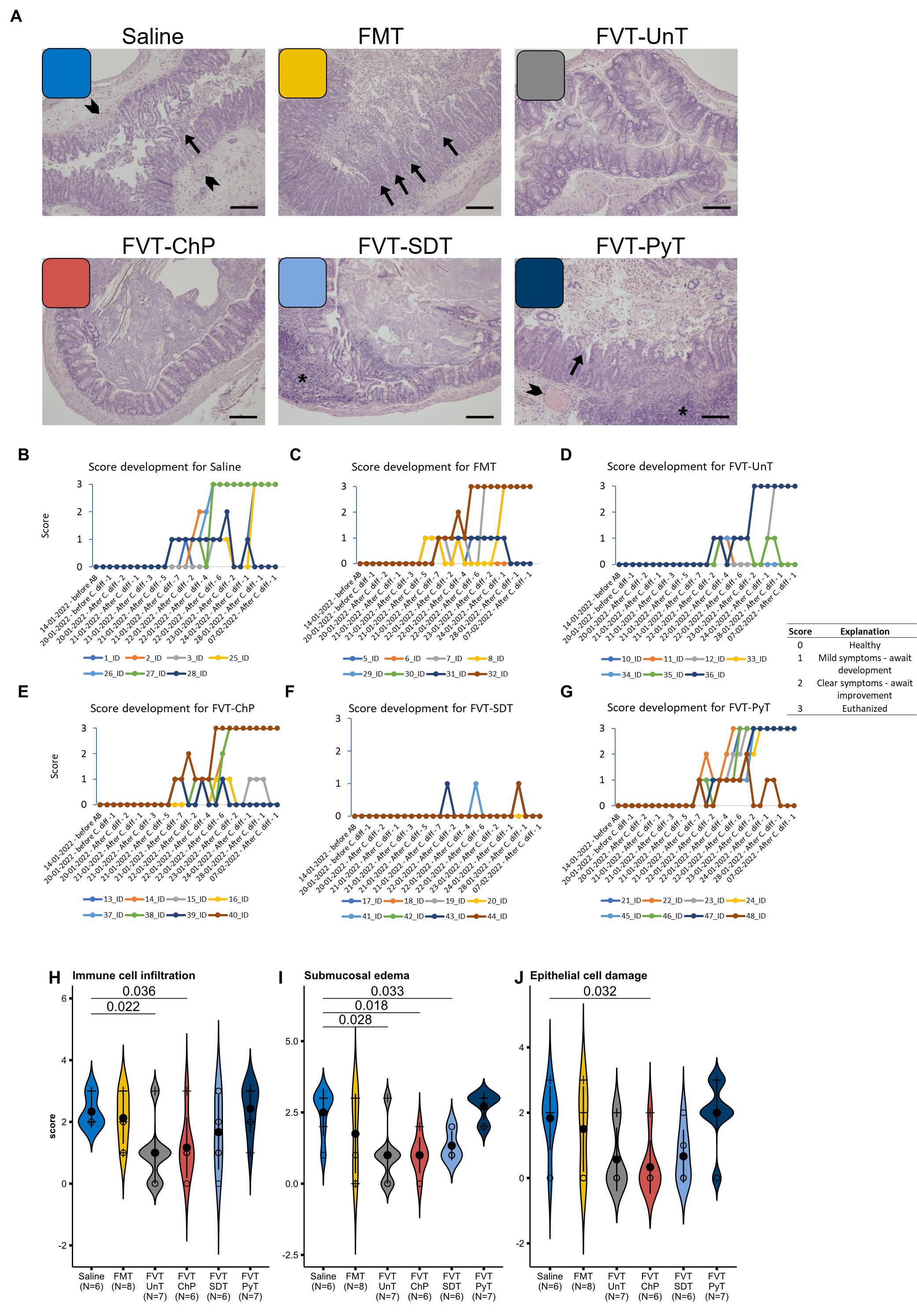

Fig. S4: Evaluation of CDI symptoms. A) Representative histology images of cecum tissue sampled at either termination of the study or when mice were euthanized due to reaching humane endpoints. Asterisks = immune cell infiltrates, arrowheads = congested submucosal blood vessels, arrows = volcano lesions contributing to pseudo-membrane formation. Histology image scale bar = 300 µm. B-F) Graphs showing the qualitative visual assessment of the health status of individual mice during the monitoring. The evaluations were based on the level of physical activity (i.e. decreased spontaneous or provoked activity), level of watery feces, body posture (hunching), and whether their fur was kept clean or not. The score of 0 (healthy), 1 (mild symptoms), 2 (clear symptoms), or 3 (mice with score 2 that did not show improvement within next checkup were euthanized). The mouse ID is indicated below each graph. H-J) Showing violin plots of each of the three parameters evaluated in the histopathology assessment of the cecum tissue. Numbers above shows the p-value.

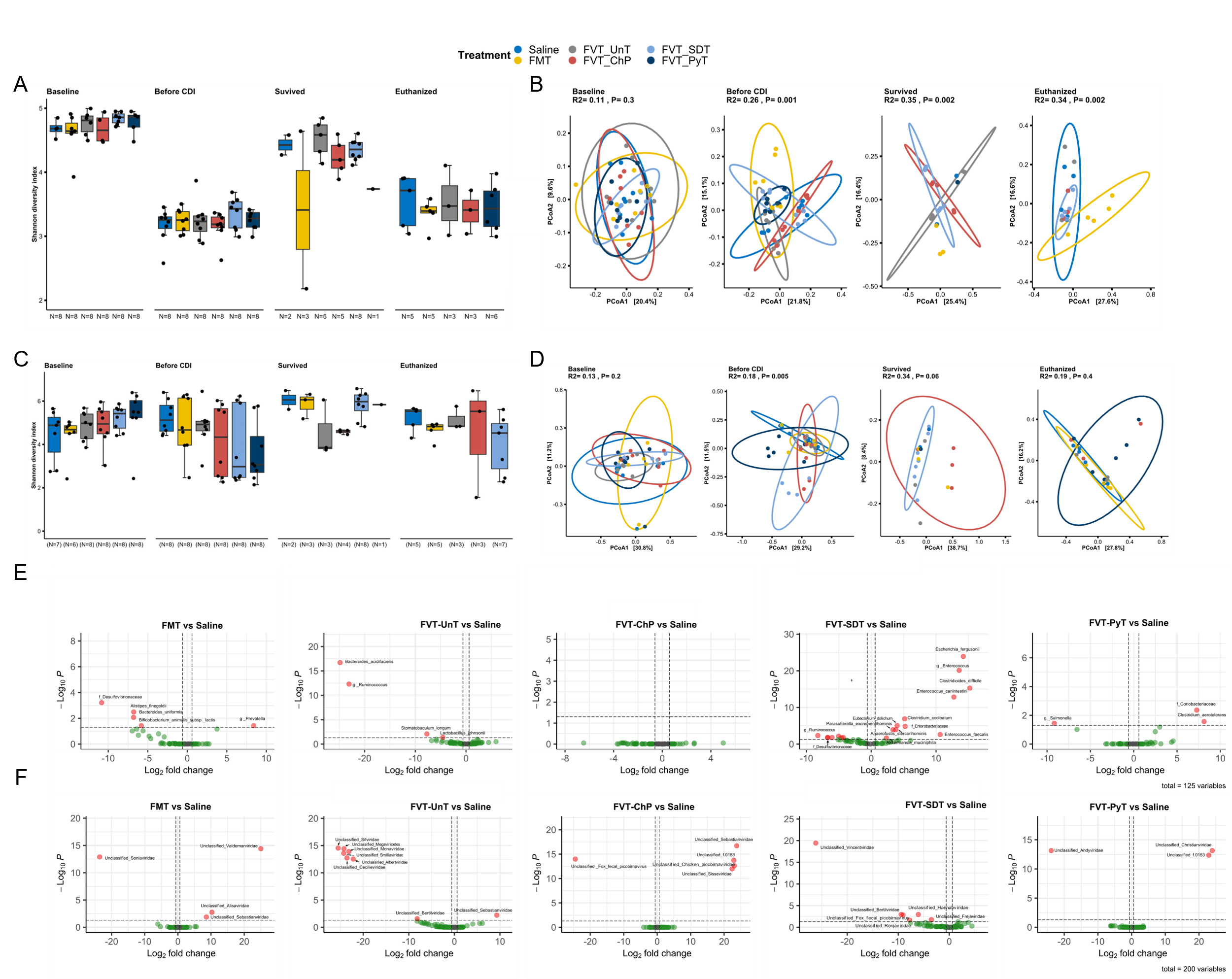

Fig. S5: Bacteriome (16S rRNA gene amplicon) and metavirome (whole-genome sequencing) analysis at three time points; Baseline (before antibiotic treatment), before CDI infection (after antibiotic treatment), and at termination (euthanized/survived). The treatment groups represent the average regardless of whether the mice survived the infection until termination or were euthanized before termination. A) & C) The bacterial and viral Shannon diversity-index (α-diversity) and B) & D) Bray-Curtis dissimilarity based PCoA plot (β-diversity). Volcano plots showing the differential relative abundance for E) the bacterial and F) viral taxa compared with the saline control.

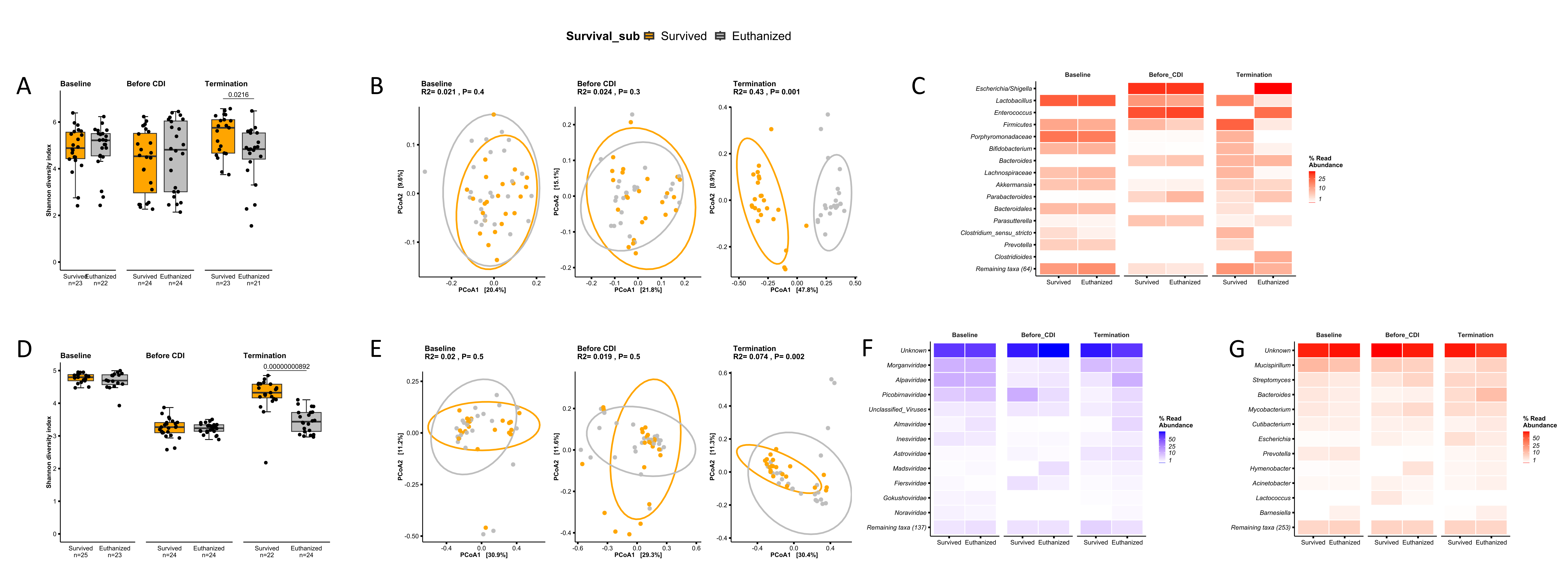

Fig. S6: Bacteriome (16S rRNA gene amplicons) and metavirome (whole-genome sequencing) analysis at three time points; Baseline (before antibiotic treatment), before *C. difficile* infection (after antibiotic treatment), and at termination of mice that either survived until termination or were euthanized regardless of the treatment. A) The bacterial Shannon diversity-index (α-diversity) and B) Bray-Curtis dissimilarity based PCoA plot (β-diversity). C) Heatmap illustrating the bacterial relative abundance in percentages of the dominating bacterial taxa that was associated to the mice that survived versus euthanized. D) The viral Shannon diversity-index and E) Bray-Curtis dissimilarity based PCoA plot. F) and G) Heatmaps illustrating the relative abundance in percentages of, respectively, the viral taxonomy and bacterial hosts that are predicted based on the viral sequences.

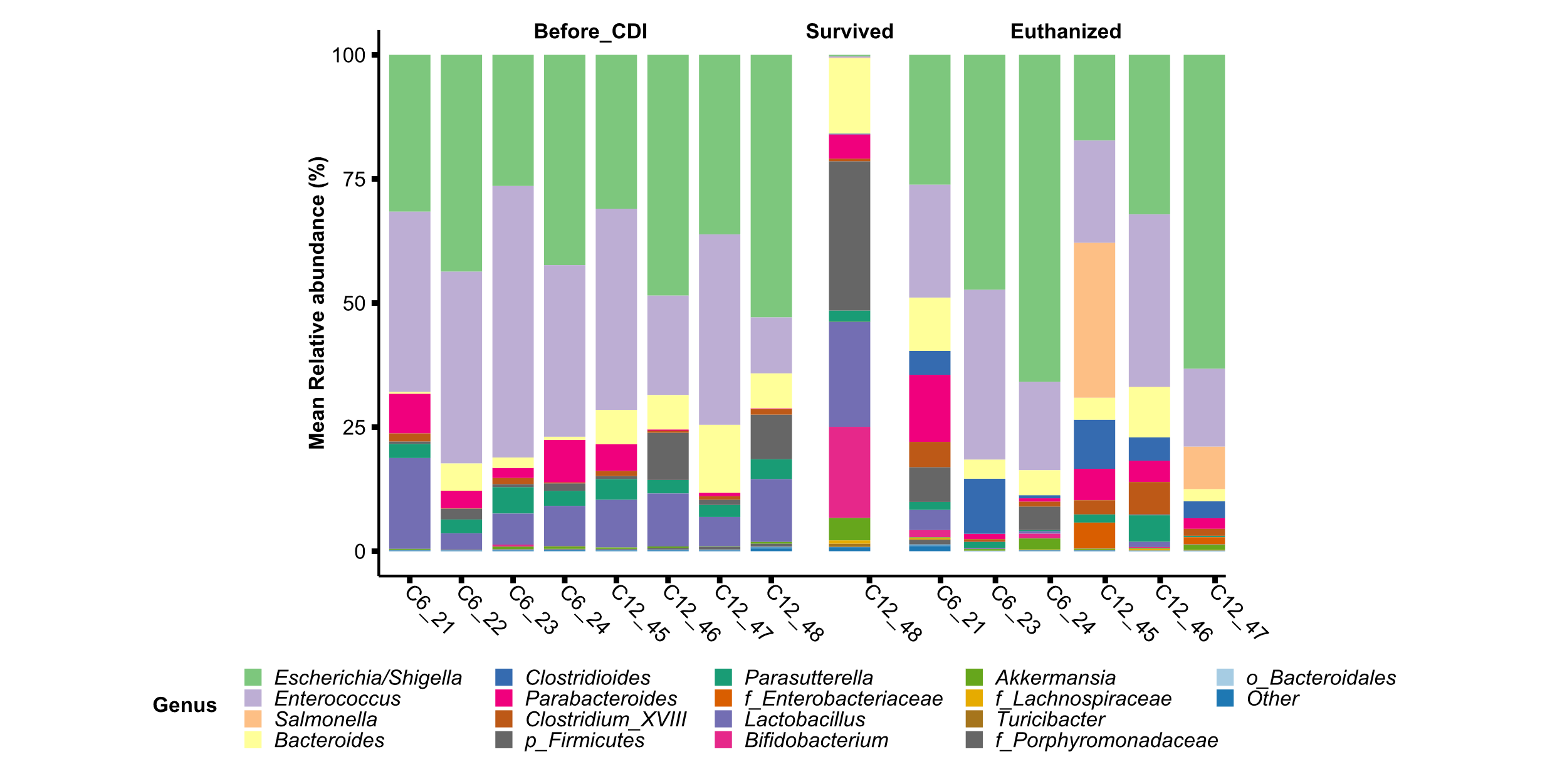

Fig. S7: Taxonomical bar plot showing the relative abundance in percentages of each individual mouse in the FVT-PyT treated group, both before the *C. difficile* infection (after antibiotic treatment) and when the mice were either euthanized or survived. Mouse No 45 and 47 had a relatively high abundance of *Salmonella* spp. that was not observed in the other mice.

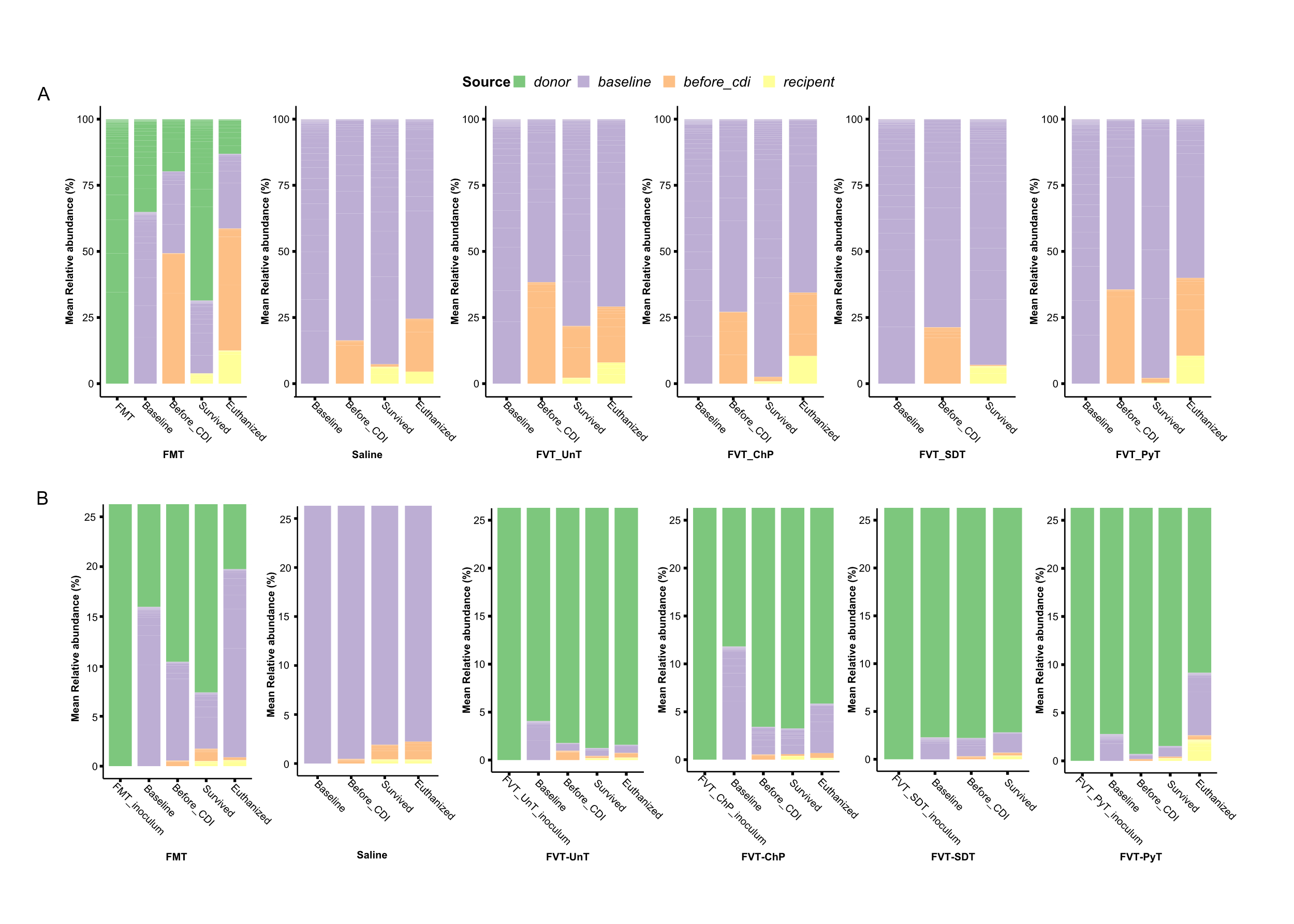

Fig. S8: The potential bacterial and viral engraftment from the FMT/FVT inoculums to the treated mice were analyzed at A) 16S rRNA gene amplicon and B) viral metagenome level (viral contigs). The gut microbiomes were divided into donor origin and the different timepoints of baseline (before antibiotic treatment), before *C. difficile* infection (after antibiotic treatment), and when the mice survived until termination or were euthanized. There was an overlap of amplicons/viral contigs found in both the FVT inoculum and in the mice at the timepoints before the treatment, hence this analysis only partly describes the bacterial and viral engraftments.

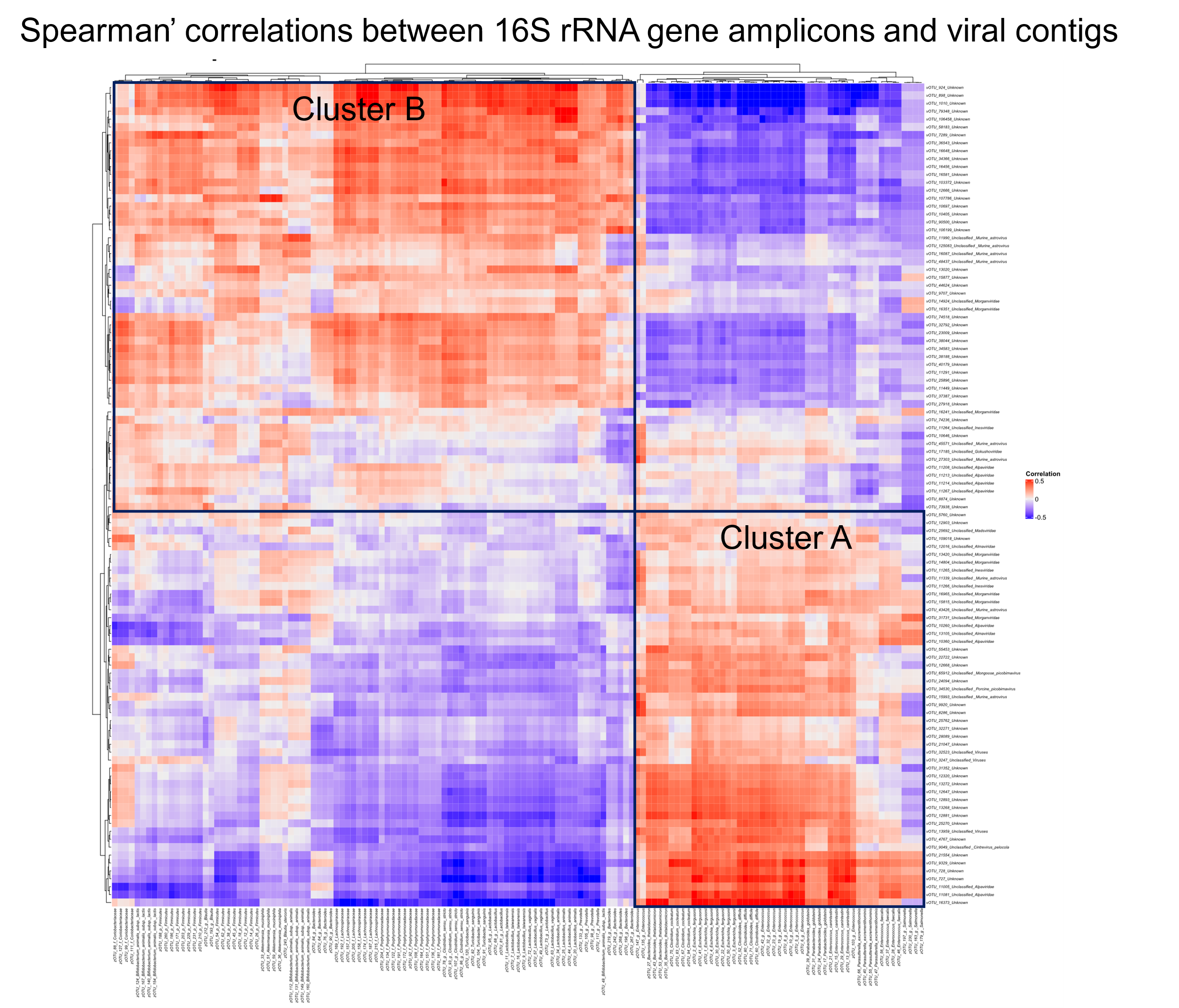

Fig. S9: Spearman’s correlation analysis of the bacterial and viral relative abundance. Cluster A and B marks bacterial and viral positive correlations.

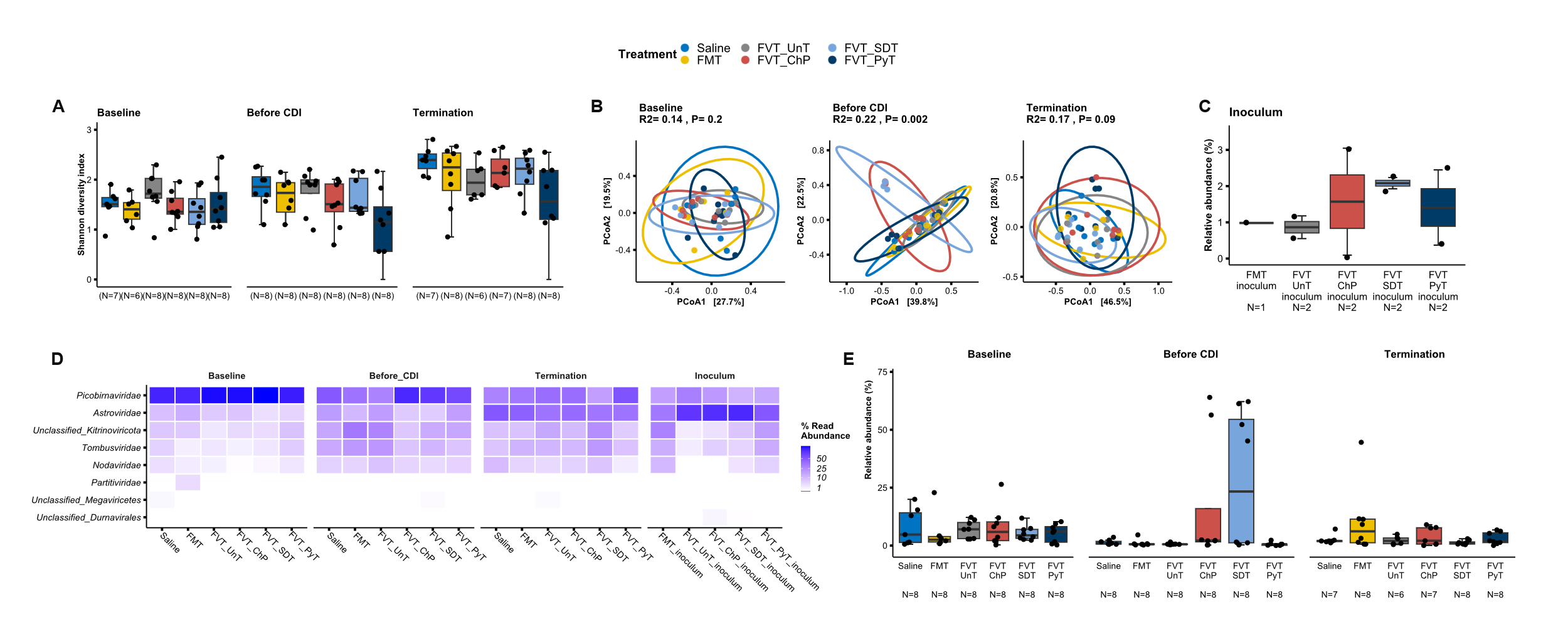

Fig. S10: Metavirome analysis of only eukaryotic viruses (that could be classified as such) based on whole-genome sequencing at three time points; baseline (before antibiotic treatment), before *C. difficile* infection (after antibiotic treatment), and at termination of mice that either survived until termination or were euthanized. A) The viral Shannon diversity-index (α-diversity) and B) Bray-Curtis dissimilarity based PCoA plot (β-diversity). C) Box plot showing the relative abundance in percentages of eukaryotic viruses in the different FMT/FVT inoculums. D) Heatmap illustrating the relative abundance of eukaryotic viral taxonomy. E) Box plot showing the relative abundance in percentages of eukaryotic viruses in the differently treated mice over time. The applied FMT and different FVTs inoculums were spiked with a defined phage mock community as a technical control of the applied protocols to detect ss/dsRNA and ss/dsDNA viruses.

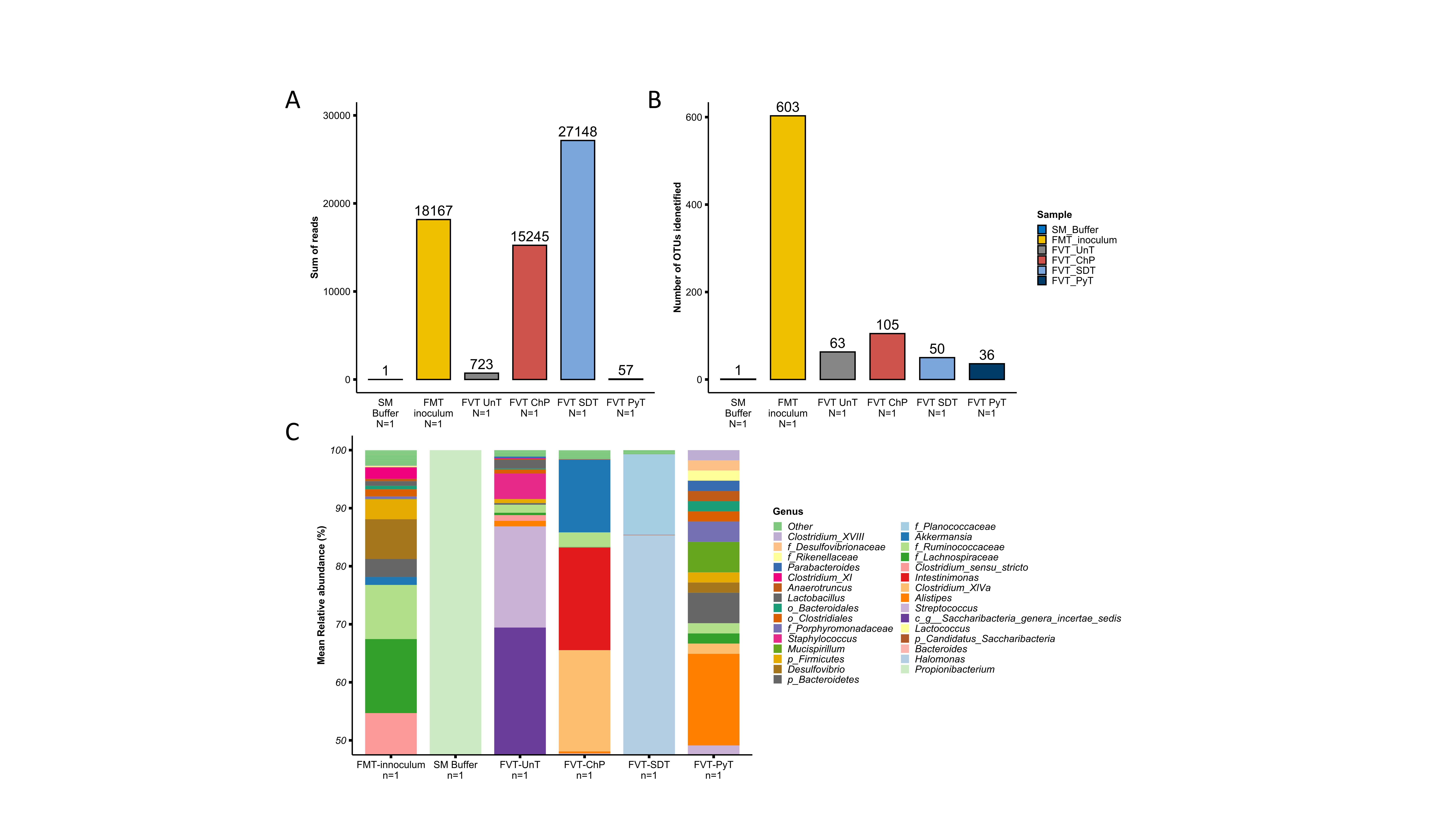

Fig. S11: 16S rRNA gene amplicon sequencing of the FMT/FVT inoculums to assess their bacterial profile. Bar plot showing A) the number of reads detected and B) the number of observed bacterial species (α-diversity), and C) the relative abundance in percentage of the bacterial taxa detected.
